# Bidirectionally connected cores in a mouse connectome: Towards extracting the brain subnetworks essential for consciousness

**DOI:** 10.1101/2021.07.12.452022

**Authors:** Jun Kitazono, Yuma Aoki, Masafumi Oizumi

**Affiliations:** Graduate School of Arts and Sciences, The University of Tokyo; Graduate School of Information Science and Technology, The University of Tokyo

## Abstract

Where in the brain consciousness resides remains unclear. It has been suggested that the subnetworks supporting consciousness should be bidirectionally (recurrently) connected because both feed-forward and feedback processing are necessary for conscious experience. Accordingly, evaluating which subnetworks are bidirectionally connected and the strength of these connections would likely aid the identification of regions essential to consciousness. Here, we propose a method for hierarchically decomposing a network into cores with different strengths of bidirectional connection, as a means of revealing the structure of the complex brain network. We applied the method to a whole-brain mouse connectome. We found that cores with strong bidirectional connections consisted of regions presumably essential to consciousness (e.g., the isocortical and thalamic regions, and claustrum) and did not include regions presumably irrelevant to consciousness (e.g., cerebellum). Contrarily, we could not find such correspondence between cores and consciousness when we applied other simple methods which ignored bidirectionality. These findings suggest that our method provides a novel insight into the relation between bidirectional brain network structures and consciousness.

## Introduction

Where in the brain consciousness resides has been one of the biggest questions in science. Although we have not yet reached a conclusive answer, much empirical evidence has been accumulated in the course of searching for the minimal mechanisms sufficient for conscious experience, called Neural Correlates of Consciousness (NCC)^1^. Among the many problems that need to be solved in identifying NCC, we focus here on the problem of identifying the minimally sufficient subnetworks in the brain which support conscious experience. In this study, we simply refer to such subnetworks as “the locus of consciousness”. For example, it is commonly agreed that the retina is not included in the locus of consciousness because it has been empirically shown that neural activities in the retina do not directly correlate with what we perceive ^2^. More importantly, a person who becomes retinally blind in adulthood continues to have vivid visual dreams^2^. As another notable example, the cerebellum is also not considered to be included in the locus of consciousness because lesions of the cerebellum do not much affect conscious experience, even though it has far more neurons than the cerebral cortex and is densely connected to the rest of the brain^3,4^. On the other hand, which cortical areas or subcortical areas are essential for consciousness are still controversial (see Boly *et al.* (2017)^5^, Odegaard *et al.* (2017)^6^, and Melloni *et al.* (2021)^7^ for general reviews, and Leopold (2012)^8^ for a review focusing on the primary visual cortex as an example).

In inferring the locus of consciousness in the brain, it is important to note suggestions that feed-forward processing alone is insufficient for subjects to consciously perceive stimuli; rather, feed-back is also necessary, indicating the need for bidirectional (also called recurrent, reciprocal, or reentrant) processing^9–17^. The feedback component disappears not only during the loss of specific contents of consciousness in awake states, but also during unconscious states, where conscious experiences are generally lost, such as general anesthesia ^10,18–20^, sleep^21^, and vegetative states^22^. The importance of bidirectional processing is suggested to be independent of sensory modality^23^ (vision ^10–12,15,16^, somato-sensation^9,13,14,17^, audition^24^) and species (humans ^13,15,19,22^, monkeys ^9–12,16,21^, rodents^14,17^, birds^25^, and even flies^20^).

Given these findings, it appears reasonable that subnetworks in which brain areas are strongly bidirectionally connected would be included in the locus of consciousness. In fact, many major theories of consciousness have made similar predictions about the locus of consciousness in common, even though they differ in many other respects^26–37^. Under this criterion, the retina, for example, is evidently excluded from the locus of consciousness because it is connected to the other areas of the brain in a purely feed-forward manner. To examine the relation between subnetworks with strong bidirectional connections and consciousness, it is important to first identify such subnetworks and understand the bidirectional network structure of the brain. If we understand which subnetworks are strongly bidirectionally connected and which are only weakly so connected, we can quantitatively discuss the correspondence between these subnetworks and consciousness.

For this purpose, we propose a method for extracting subnetworks in which nodes are strongly connected in a bidirectional manner. We call such subnetworks “complexes”; this term and concept originate from integrated information theory, although the specific definition of complexes differs in the original theory^31,33,38–41^. To be specific, in this study, we first define a “main” complex as a subnetwork that has the local maximum of bidirectional connection strength. We evaluate the strength of bidirectional connections by a measure that takes a large value when nodes are connected by strong bidirectional edges. To reveal the network structure, we also define complexes as a weaker notion of a main complex. Complexes are less strongly bidirectionally connected than a main complex and form a nested structure. That is, a main complex is included in another less strongly connected complex; that complex is in turn included in yet another complex; and so on. In this hierarchical organization, a main complex, intuitively speaking, is a central core where there is no weakly connected parts and complexes are surrounding cores.

If we search for complexes by brute force, computation time grows exponentially with the number of nodes, because we need to take account of all possible subnetworks. To reduce computation time, we can use an algorithm proposed in our previous study^41^. This algorithm, called hierarchical partitioning for complex search (HPC), enables the identification of complexes simply by hierarchically dividing the entire network. Because of the simplicity of this algorithm, the computation time increases only polynomially with the number of nodes. HPC allows us to find all complexes in a practical amount of time, even from large networks of thousands of elements, without omissions or misidentifications.

As a step in investigating the relationship between bidirectionally connected subnetworks - complexes - and consciousness, we applied the proposed method to a meso-scale, whole-brain mouse connectome^42^ and identified the complexes. This connectome includes not only the cortical regions but also subcortical, brainstem and cerebellar regions, and has high spatial resolution. These characteristics make it suitable for discussing the relationship between brain regions and consciousness. We found that the extracted complexes with strong bidirectional connections consist of the brain regions that are thought to be essential to consciousness. In addition, to assess whether it is important to take account of the bidirectionality of connections, we examined how the results are affected if the bidirectionality of connections is ignored. We found that the complexes do not necessarily consist of the particular brain regions thought to be essential to consciousness, but rather of various brain regions that do not directly contribute to consciousness. We also applied a widely used method for extracting network cores, *s*-core decomposition, which does not consider bidirectionality. Interestingly, we could not find such correspondence between the extracted cores and the brain regions presumably essential to consciousness. In addition, we investigated the relationship between the complexes and the degree of nodes. We found that the complexes with strong bidirectional connections do not necessarily consist of high-degree nodes. This means that the core structures revealed by the complexes largely differ to the structures that are revealed by degree-based methods that ignore bidirectionality. These results indicate that the identification of bidirectional network structures will provide new insights into areas essential to consciousness.

## Results

### Network cores with strong bidirectional connections: Complexes

#### A simple example of a complex

In this study, we tried to extract the bidirectionally connected “cores” of the network, called “complexes”. Before we introduce the definition of complexes, let us first intuitively explain the concept of complexes taking the network shown in Fig. 1 as an example. In this example, the node *A* and the nodes *CDGH* are only unidirectionally connected to *BEFIJ*, and therefore these nodes are not included in a complex. The node set *BEFIJ* is a complex but only a weakly connected one, because the node *B* is only weakly connected to the nodes *EFIJ* (i.e., there is only one edge in each direction, *B* to *EFIJ* and *EFIJ* to B). In contrast, the nodes *EFIJ* are all strongly connected to each other, and the nodes *EFIJ* therefore constitute a strongly connected complex. In this example, the node set *EFIJ* turns out to be the most strongly connected complex, which we call the main complex. In general, complexes form a nested hierarchical structure, as do the node sets *EFIJ* and *BEFIJ*. That is, a complex contains another complex that is smaller in size but more strongly connected. The complex smallest in size is the most strongly connected, and thus the main complex.

**Figure 1:**
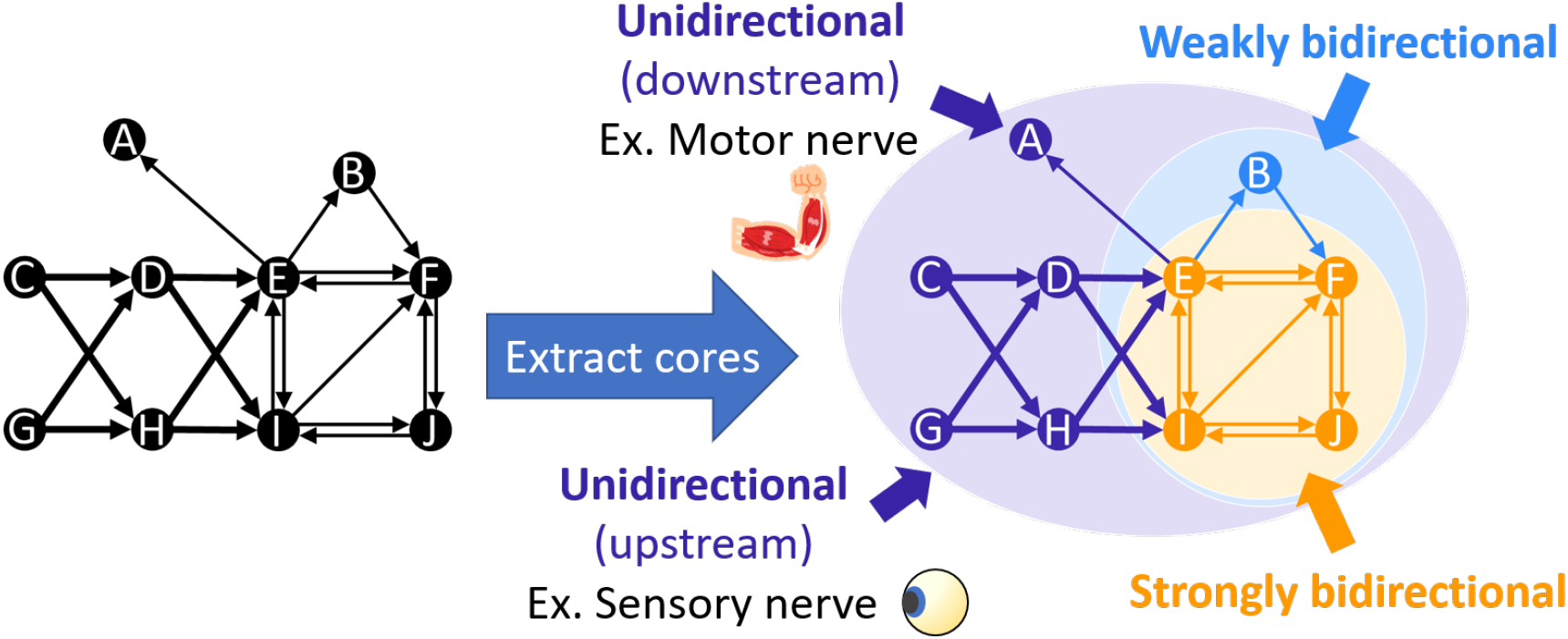
Schematic of extracting network cores complexes and main complexes. A complex is a subnetwork that consists of stronger bidirectional connections than other subnetworks that include it. If we extract complexes from the network on the left, we obtain the result on the right. There are two complexes. One is the subnetwork *EFIJ*, which is colored orange. The other is the subnetwork *BEFIJ*, which is colored light blue. Comparing the two subnetworks, *EFIJ* is more strongly bidirectionally connected than *BEFIJ* because the nodes *EFIJ* are all strongly connected to each other while the node *B* is only weakly connected to the nodes *EFIJ* (i.e., there is only one edge in each direction, *B*to *EFIJ* and *EFIJ* to *B*). The node set *EFIJ* turns out to be the most strongly connected complex in this network, which we call the main complex. The entire network is not a complex. The strength of the bidirectional connections in the entire network is zero because the entire network includes the nodes *CDGH* and *A*, which are connected in a completely feed-forward manner. If we compare this network to the nervous system of the whole body of a mammal, we can consider the bidirectionally connected nodes *BEFIJ* as the brain, the nodes *CDGH* upstream as sensory nerves such as the retina, and the downstream node *A* as motor nerves.

We can consider this exemplar network as a toy network of the nervous system. For instance, we can consider the bidirectionally connected nodes *BEFIJ* as the brain, the upstream nodes *CDGH* as sensory nerves such as the retina (afferent nerves), and the downstream node *A* as motor nerves (efferent nerves). As we explain above, the node *A* (motor nerves for example) and the nodes *CDGH* (the retina for example) are not included in the complexes. If we assume that bidirectional processing is essential for consciousness, the motor nerves and the retina would not be included in the locus of consciousness. In the mouse connectome network investigated in this study, there are no nodes that are only unidirectionally connected to the rest of the network. Thus, we cannot evidently exclude some of the nodes because of the lack of bidirectional connections. Rather, we need to quantitatively investigate the degree of the bidirectional connections and look at the hierarchical structure of the complexes.

#### Outline of complexes and related concepts

The mathematical definition of a “complex” is rather complicated. To get the gist of it, we first outline two important concepts, namely strength of bidirectional connections and minimum cut, and then outline complexes. Please see Methods for mathematically formal explanations.

##### Strength of bidirectional connections

To define complexes, i.e. bidirectionally connected cores of a network, we first need to have a measure that quantifies how strongly the two divided parts of a network are bidirectionally connected. We propose a measure that is low when the connections in one direction are weak even though those in the other direction are strong (Figs. 2a and 2b) and that is high when the connections in both directions are strong (Fig. 2c). Specifically, we define the strength of bidirectional connections as the minimum value of the sum of the weights of connections going from one part to the other and the sum in the opposite direction (Eq. (6) in Methods). The strength of the bidirectional connections defined this way is zero when the connections are completely unidirectional as in Fig. 2a, low when the connections in one direction are weak as in Fig. 2b, and high when the connections are strong in both directions as in Fig. 2c.

**Figure 2:**
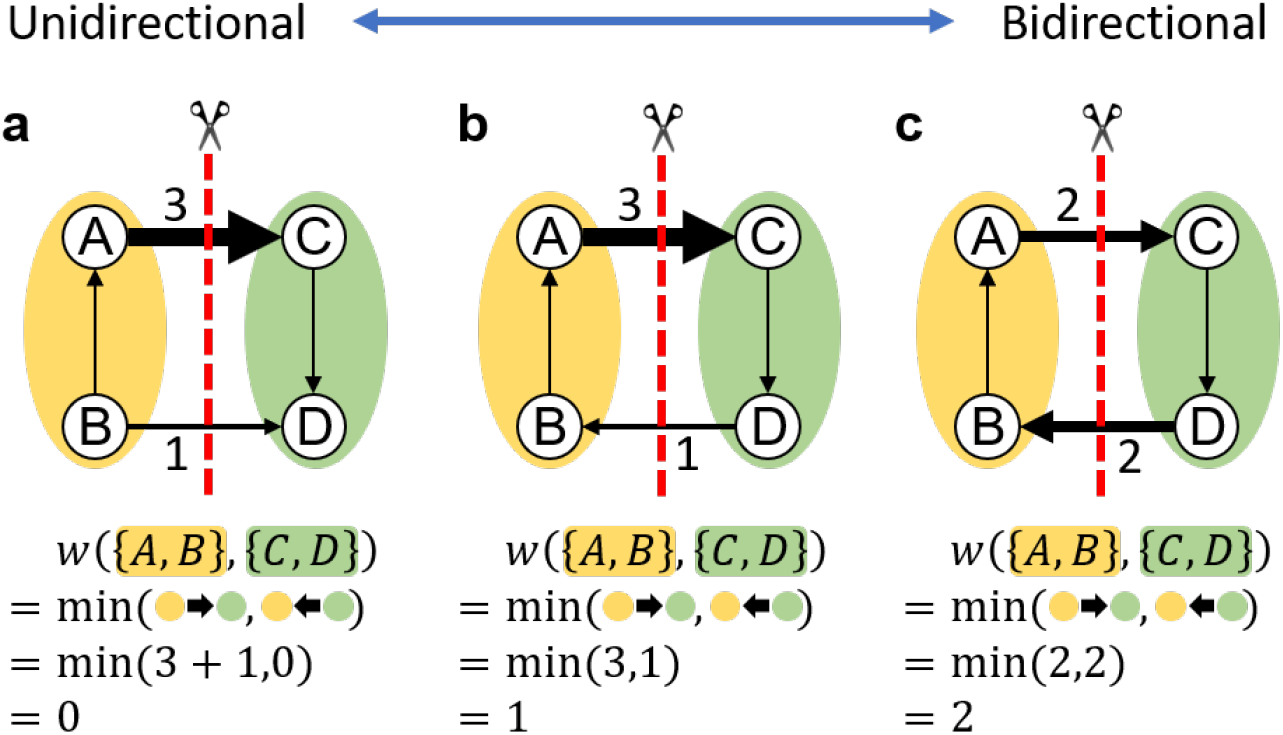
Strength of bidirectional connections. To measure the strength of bidirectional connections, we take the minimum value of the sum of the weights of connections going from one part to the other and the sum in the opposite direction. In the examples **a–c**, the strength of bidirectional connections between *AB* and *CD* is measured. **a** The connection is completely unidirectional: there are connections from *AB* to *CD* but there are no connections in the opposite direction. In this case, the strength of bidirectional connections is 0. **b** The connection from *AB* to *CD* is strong but that from *CD* to *AB* is weak. In this case, the strength of bidirectional connections is low, which equals to 1. **c** The connections in both directions are strong and the strength of bidirectional connections is high, which equals to 2.

##### Minimum cut

A complex is a network core whose parts are strongly connected to each other in a bidirectional manner. In other words, a complex cannot be “cut” into two parts without losing many strong edges no matter how it is cut. To measure such “inseparability” of a network, we consider the bi-partition of the network for which the strength of bidirectional connections is minimum among those for all possible bi-partitions, which we call a minimum cut (or a min-cut). We call the strength of bidirectional connections for a min-cut the “min-cut weight” and represent it by *w*^mc^. As the value of a min-cut weight *w*^mc^ gives the lower bound of the strength of bidirectional connections for any possible bi-partitions of the network, any part of the network is “bidirectionally” connected to its complement part with a strength greater than or equal to *w*^mc^.

If a network consists of disconnected parts as shown in Fig. 3a, the min-cut is the partition that cuts the network into the two disconnected parts and *w*^mc^ is 0. On the other hand, in a network where all the parts are strongly connected and cannot be separated without many edges being cut as in Fig. 3b, *w*^mc^ is large. As illustrated in these examples, a larger min-cut weight *w*^mc^ indicates a network that is more inseparable.

**Figure 3:**
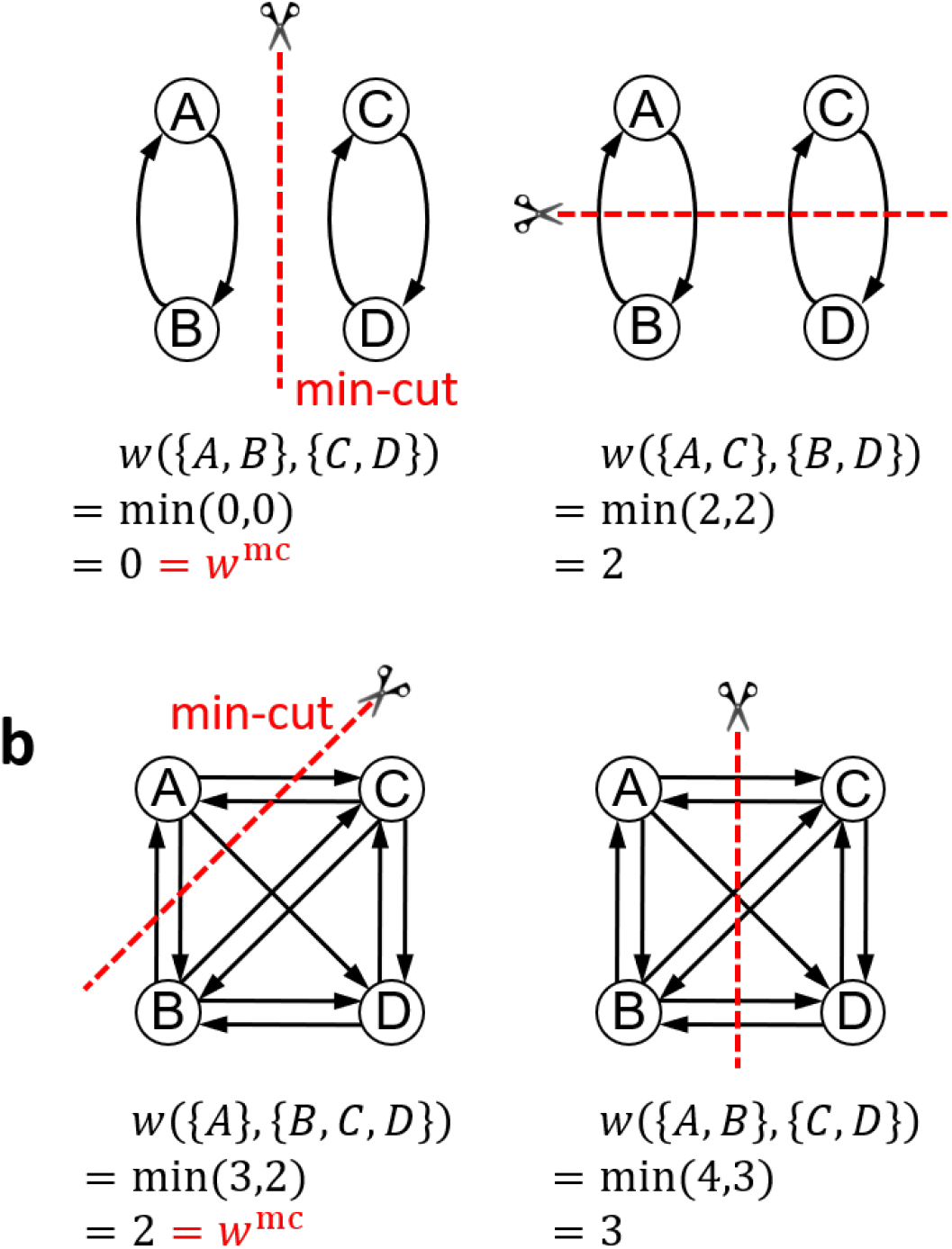
Schematic of minimum cut. The minimum cut (min-cut) is the cut for which the strength of bidirectional connections is minimum among all possible cuts. In this figure, we assume that all edge weights are 1. **a** A network consisting of two mutually disconnected groups *AB* and *CD*. The min-cut partitions this network into the two parts *AB* and *CD*, and its weight is zero (*w*^mc^ = 0). On the other hand, if the network is cut into *AC* and *BD*, the strength of bidirectional connections becomes nonzero. **b** A network where all the parts are strongly connected in a bidirectional manner. This network cannot be separated without cutting many edges. The strength of bidirectional connections is therefore high, even for its min-cut (*w*^mc^ = 2).

##### Complex

Complexes and main complexes are defined using the min-cut weight *w*^mc^ introduced above. A main complex is a subnetwork that has “locally” maximal *w*^mc^. Local maximum means that *w*^mc^ in a main complex is larger than that in any other subnetwork containing it and any other smaller subnetwork contained within it (both the left and right inequality in Fig. 4 hold). In general, a network can have multiple main complexes. In addition to main complexes, the notion of complexes is also useful for revealing the structure of a network. Briefly, a complex is a weaker notion of a main complex, i.e., a subnetwork such that its *w*^mc^ is greater than *w*^mc^ of any other subnetwork containing it (only the right inequality in Fig. 4 holds). Complexes form a hierarchical structure: a main complex is included in a complex larger in size but with smaller *w*^mc^, and the complex is included in yet another complex even larger in size but with smaller *w*^mc^. Metaphorically speaking, if we consider a network as a mountain whose height is determined by the min-cut weight *w*^mc^, a main complex tells us the peak of the mountain and the surrounding complexes tell us the contour lines of the mountain, as illustrated in Fig. 1.

**Figure 4:**
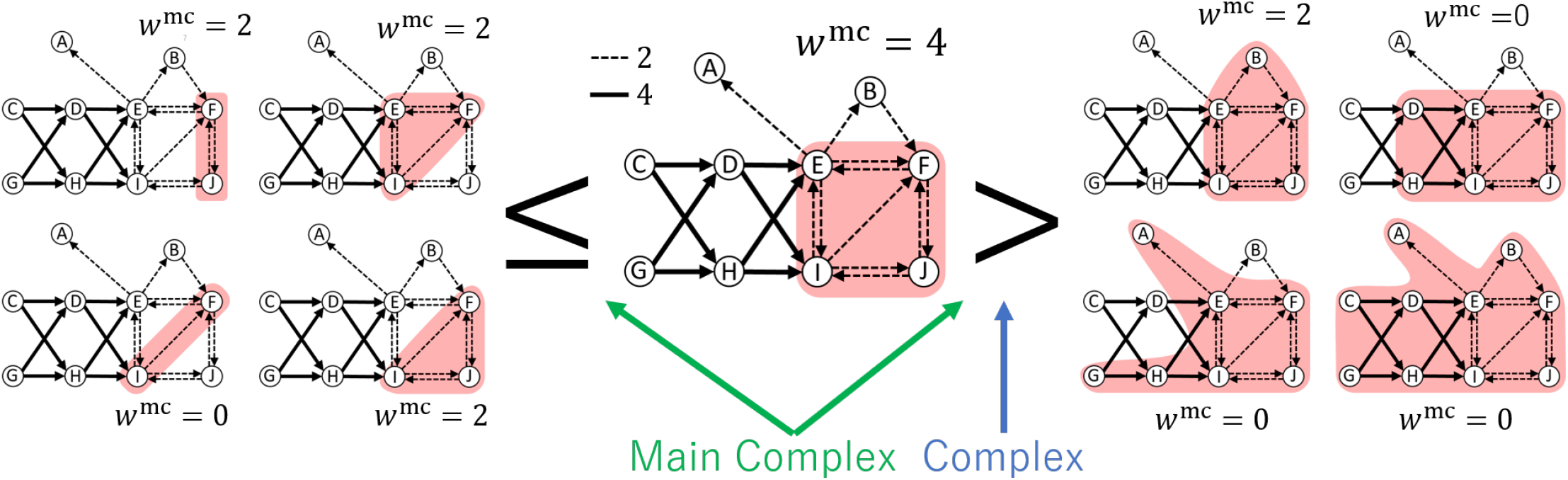
Schematic of the definition of complexes and main complexes. The subnetwork {*E,F,I,J*} is a complex because it has a greater min-cut weight *w*^mc^ than any larger subnetworks that contain it, namely, {*B,E,F,I,J*}, {*D, H, E, F, I, J*}, {*A, E, F, G, H, I, J*}, and so on. In addition, the subnetwork {*E, F, I, J*} is a main complex because it has a greater min-cut weight than not only the larger subnetworks but also any smaller subnetworks within it, namely, {*E, J*}, {*F, I*}, {*E, F, I*}, and so on.

A schematic explanation of complexes is shown in Fig. 4. We consider a network, which is the same as that in Fig. 1. For example, the subnetwork consisting of the four nodes {*E,F,I,J*} is a complex because its min-cut weight *w*^mc^ is greater than any larger subnetworks containing it ({*B, E, F, I, J*}, {*D, E, F, H, I, J*}, etc.). In addition, the subnetwork {*E, F,I,J*} is also a main complex because its *w*^mc^ is greater than those of not only larger subnetworks but also of any smaller subnetworks contained within it ({*F, J*}, *{E,F,I*}, etc.).

### Extracting complexes

#### Fast and exact algorithm to search for complexes

If complexes are searched for by brute force, the computation time increases exponentially with the number of nodes *N*. This is because it is necessary to compute the min-cut weight *w*^mc^ for all of the *O*(2^*N*^) subnetworks and then compare these values. On the other hand, by using a fast and exact method we proposed in our previous study, Hierarchical Partitioning for Complex Search (HPC)^41^, we need to compute *w*^mc^ for only *N* – 1 subnetworks. As a result, the overall computation time increases only polynomially with *N*, and it is possible to analyze networks with several thousands of nodes in a practical time (See Supplementary Fig. 1 for an actual computation time evaluated by a simulation).

We illustrate how HPC works using the example shown in Fig. 5. In the following, for simplicity of notation, we write a subnetwork consisting of a node set *S* simply as *S*. In HPC, a network is hierarchically partitioned by min-cuts until it is decomposed into single nodes. First, the whole network *V* = {*A, B, C, D, E, F, G*} is divided by its min-cut (indicated by a dashed line) into *V*_L_ = {*A,B,E,F*} and *V*_R_ = {*C,D,G*}. Then, *V*_L_ is divided into *V*_LL_ and *V*_LR_, and *V*_R_ into *V*_RL_ and {*G*}. Finally, the whole network *V* is decomposed into seven single nodes. After this process, we obtain the set of hierarchically partitioned subnetworks *V*, *V*_L_, *V*_R_, *V*_LL_, *V*_LR_, *V*_RL_. We consider all the set of subnetworks 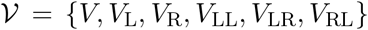, excluding single nodes. We can then mathematically prove that any complex in the network belongs to 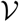. The proof is based on the mathematical property “monotonicity” and its satisfaction by the strength of bidirectional connections (Eq. (6) in Methods). Thus, we can consider 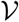 as the candidate complexes. We can select complexes from these 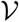 candidates without omissions or misidentifications. See Methods for more details.

**Figure 5:**
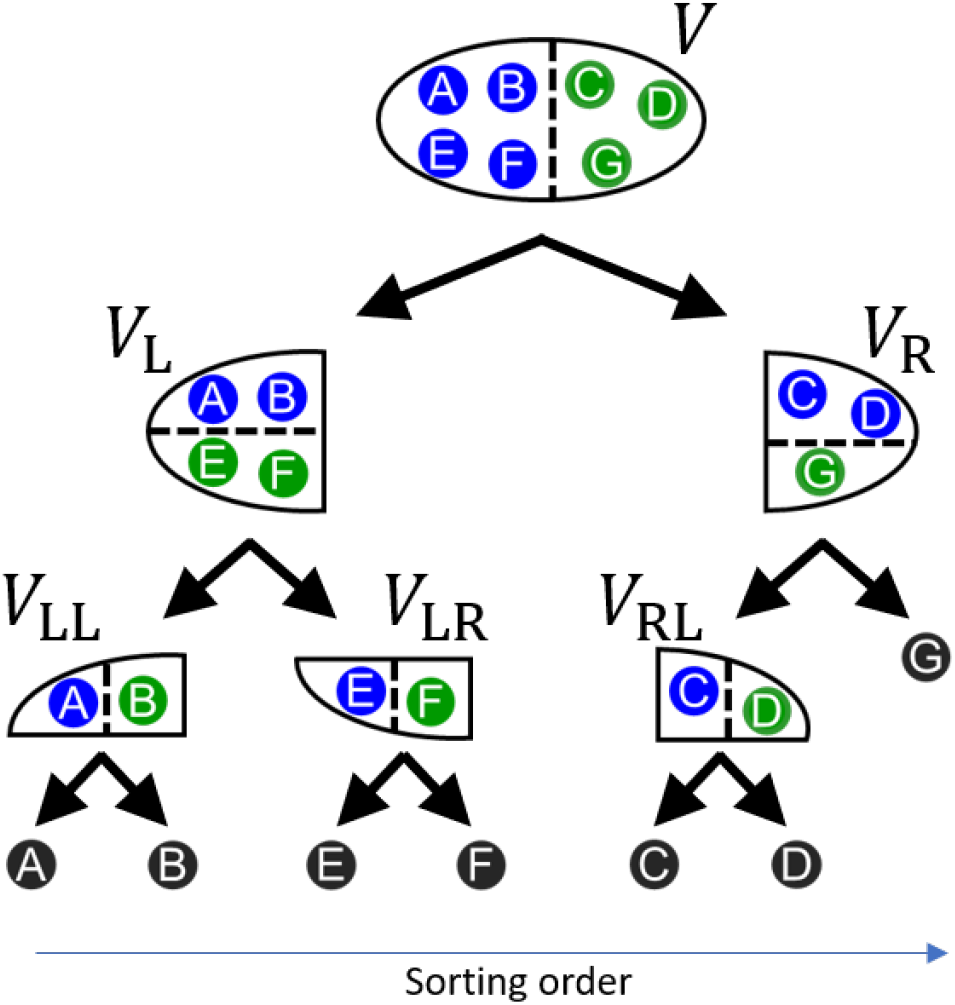
Schematic of Hierarchical Partitioning for Complex search (HPC). In HPC, a network is hierarchically partitioned by min-cuts until the network is decomposed into single nodes. In this example, the whole network *V* = {A, *B, C, D, E, F, G*} is divided by its min-cut (indicated by a dashed line) into *V*_L_ ={*A, B, E, F*} and *V*_R_ = {*C, D, G*}. Then, *V*_L_ is divided into *V*_LL_ and *V*_LR_, and *V*_R_ into *V*_RL_ and {*G*}. Finally, the whole network *V* is decomposed into seven single nodes. In this process, we only need to evaluate the *w*^mc^ of *N* – 1 (=6) subnetworks. This number is much smaller than the number of subnetworks evaluated in the brute force method, 2^*N*^ – *N* – 1 (= 57), which is the number of subnetworks consisting of more than one node. The subnetworks appearing in this hierarchical partitioning process are candidate complexes. The bottom arrow indicates how we sort rows (columns) of the connection matrices in Figs. 6d and 6h and 7b and 7e (See Methods).

In this process, we need to evaluate *w*^mc^ of only *N* – 1 (= 6) subnetworks, which are the subnetworks in 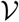. This number is much smaller than the number of subnetworks evaluated in the brute force method, 2^*N*^ – *N* – 1 (= 57), which is the number of subnetworks consisting of more than one node.

#### Complexes in a network form a hierarchical structure

As we mention above, we can find complexes from among the candidate subnetworks 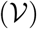 appearing in the hierarchical partitioning process. Since the candidate subnetworks form a nested hierarchical structure, as we can see in Fig. 5, complexes in a network consequently form a nested hierarchical structure as in Fig. 1. That is, a complex contains another complex that is smaller in size but has a greater *w*^mc^. A complex that is locally the smallest in size has a locally maximum *w*^mc^, which is a main complex. See Methods for mathematical details.

Please note that a nested hierarchical structure is not necessarily a single peak structure, but can have multiple peaks (i.e. there can be multiple main complexes in a network). For example, in Supplementary Fig. 2, there are two main complexes and the complexes form a nested hierarchical structure with the two main complexes as peaks.

### Demonstration of the proposed method in a toy example

In this subsection, we demonstrate with a simple example how we can understand the structure of a network by extracting the complexes. We will also explain how to visualize the results, which will be used in showing the results of the mouse connectome analysis in the next subsection. In addition, to show the significance of considering bidirectionality, we illustrate using the same example how the results are affected if bidirectionality is not considered. Finally, to highlight the characteristics of the proposed method, we compare it with a representative method for extracting network cores, s-core decomposition, which does not consider the bidirectionality of connections.

#### Understanding a network structure based on complexes

We consider the network shown in Figure 6a, which is the same as that in Fig. 1. We visualize the complexes in this network in Fig. 6b. As mentioned in the description of Fig. 1, there are two complexes in this network. One is the node set {*E,F,I,J*} (indicated by orange), and the other is the node set {*B,E,F,I, J*} (indicated by light blue), and their min-cut weight *w*^mc^ values are 2 and 1, respectively. The node set *EFIJ* is the main complex. The whole network with non-zero *w*^mc^ is always a complex because it is not contained in a larger subnetwork. However, in this case, the min-cut weight *w*^mc^ of the entire network is 0. Thus, the entire network is not a complex because we do not call a network a complex when its *w*^mc^ is 0, i.e., it is completely separable. From this figure, we can see that the two complexes are nested. That is, the complex {*E,F,I,J*}, which has a larger *w*^mc^, is contained in {*B,E,F,I, J*}, which has a smaller *w*^mc^.

**Figure 6:**
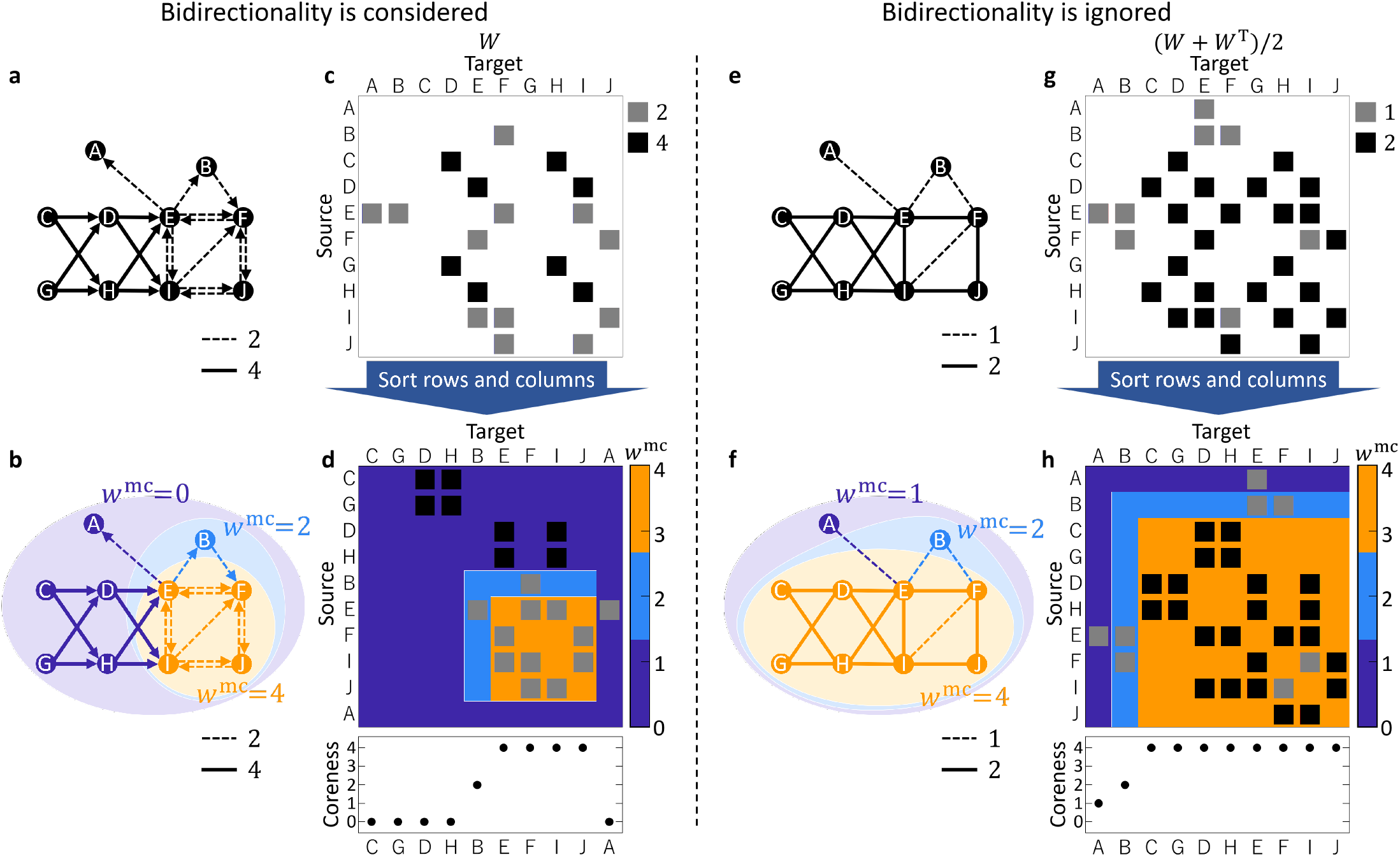
Complexes in a toy network. Bidirectionality is considered in **a–d** and ignored in **e–h**. **a** A toy network, which is the same as the network in Fig. 1. **b** The structure of complexes. Each complex is indicated by a color representing the min-cut weight *w*^mc^. **c** The connection matrix of the network in **a**. Edge weight is shown in grayscale: white, gray and black indicate 0, 1 and 2, respectively. **d** The rows and columns of the connection matrix are sorted according to the hierarchical structure of the complexes. The color map changing from blue to orange indicates the min-cut weight *w*^mc^ of the complexes. Square areas correspond to the complexes and they are superimposed in the ascending order of *w*^mc^. The plot at the bottom shows coreness values. **e** The same network as in **a** except that the direction of connections are ignored, which corresponds to ignoring bidirectionality of connections (see Methods). **f–h** The connection matrix, the complexes, and the sorted connection matrix and the coreness as in **b–d**.

We can also visualize complexes using connection matrices (Figs. 6c and 6d). In Fig. 6d, the rows and columns of the connection matrix (Fig. 6c) are sorted according to the hierarchical structure of the complexes (See Methods for a detailed description of the sorting process). In Fig. 6d, the color map indicates the min-cut weight *w*^mc^ of the complexes. Square areas correspond to the complexes and are superimposed in ascending order of *w*^mc^. We can see that the colored square areas in the sorted connection matrix in Fig. 6d corresponds to the colored areas in Fig. 6b.

By using the complexes and their *w*^mc^, we can also see how each node is distributed in complexes with different strength (*w*^mc^). For example, node *E* is included in the strongest complex, i.e. the complex with the highest *w*^mc^, and node *B* is included in the weaker complex, and so on. To quantify the strength of the complexes that each node is included in, we use an index called “coreness”. We define the coreness of node *v* as *k_v_* if node *v* is included in a complex with *w*^mc^ = *k_v_* but not included in any complex with *w*^mc^ > *k_v_* (Eq. (11) in Methods). The coreness values correspond to the color of the nodes in Fig. 6b, and in the same way, to the color of the diagonal elements in Fig. 6d. From this figure, we can see, for example, that nodes *E*, *F*, *I*, and *J* have the largest value of coreness, which means that they are included in a complex with the largest value of *w*^mc^. On the other hand, the nodes *A*, *C*, *D*, *G*, and *H* have a value of 0 for coreness, indicating that they are not included in a complex with *w*^mc^ > 0.

#### Effect of considering bidirectionality

To illustrate the significance of considering bidirectionality, we compare the complexes extracted when considering bidirectionality with those extracted when ignoring bidirectionality. When we ignore bidirectionality, we quantify the strength of connections by the sum of all the edge weights between two parts (divided by 2 for consistency with the case when considering bidirectionality) as in Eq. (3) in Methods. Quantifying the strength of connections with this simple measure is equivalent to quantifying the strength of bidirectional connections with the original measure (Eq. (6) in Methods) after symmetrizing a network (i.e. taking the mean of the original connection matrix *W* and its transpose *W^T^*) to make it virtually undirected. See Methods for more details.

Figure 6e represents the same network as that in Fig. 6a but the direction of the connection is ignored. The symmetrized connection matrix (*W* + *W*^T^)/2 is shown in Fig. 6g. Figures 6f and 6h show the results of the extracted complexes in this undirected network. Unlike the case when bidirectionality is considered (Figs. 6c and d), the main complex contains not only *E,F,I*, and *J* but also *C, D, G*, and *H*. This is because the nodes *C*, *D*, *G*, and *H* are strongly unidirectionally connected to other nodes but not bidirectionally connected. Also, *w*^mc^ for the entire network is nonzero but is zero in the original directed network. Reflecting the structure of the complexes, the coreness values are highest for the nodes *C*, *D*, *E*, *F*, *G*, *H*, *I*, and *J*, and the coreness of every node is nonzero (Fig. 6h). As can be seen in this example, if the bidirectionality of connections is ignored, i.e., only the summed strength of connections is considered, the structure of the complexes is substantially changed.

A representative existing method for extracting cores of a network is *s*-core decomposition^43–46^, which does not consider the bidirectionality of connections (Supplementary Text 1). When *s*-core decomposition is applied to the network in Fig. 6, the obtained *s*-cores are identical to the complexes when bidirectionality is ignored. In general, it can be mathematically proven that *s*-cores are identical to complexes when bidirectionality is ignored under a certain condition (see Supplementary Text 1 for details). In this example, the condition holds, and accordingly the extracted *s*-cores and the complexes are exactly the same.

#### Complexes in a mouse connectome

To demonstrate whether our method is able to extract meaningful bidirectionally connected cores in a brain network, we applied it to a mouse connectome^42^ and extracted complexes. We consider this mouse connectome to be highly suitable for this purpose because it includes not only the cortical regions but also subcortical, brainstem, and cerebellar regions, and has high spatial resolution.

It consists of 213 brain regions in each hemisphere, giving 426 nodes in total. Each brain region is at a mid-ontology level and is classified into one of the major brain regions such as the isocortex, thalamus, and cerebellar cortex. The connection matrix is shown in Fig. 7a. The color coding at the top and left of the connection matrix indicates the major brain regions. The color of each entry in the matrix indicates the edge weight between the brain regions that can be considered to be proportional to the total number of axonal fibers projecting from one region to the other. See Oh *et al.*^42^ for a detailed description.

**Figure 7:**
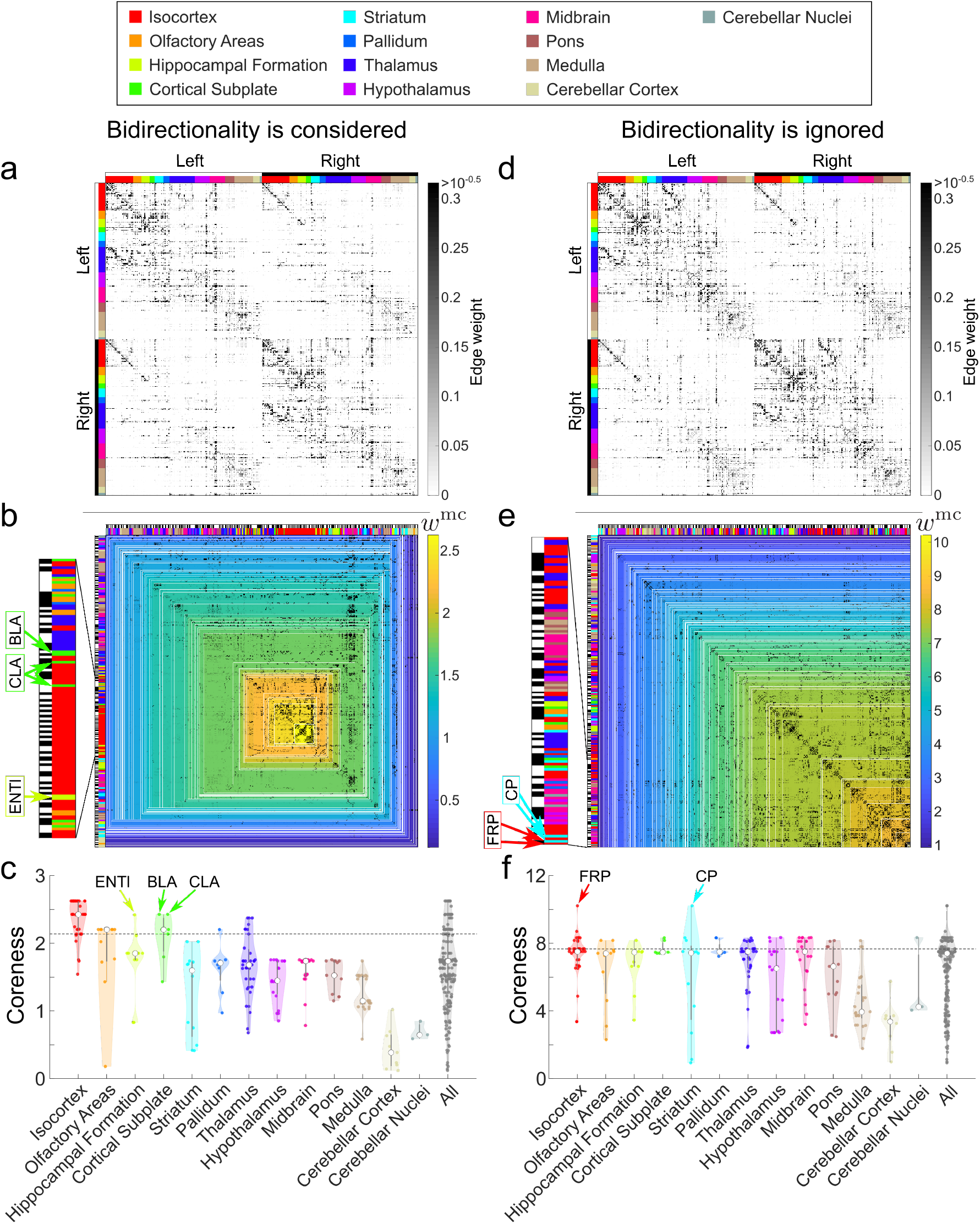
Complexes in a mouse connectome. Bidirectionality is considered in panels **a–c** and ignored in panels **d–f**. **a** The inter-region connection matrix of the mouse connectome. The color bars at the left and top of the matrix represent major brain regions and whether they are in the left or right brain. **b** Structure of complexes. A connection matrix in which the rows and columns are sorted according to the hierarchical structure of complexes as in Fig.6. The change in color map from blue to yellow indicates the min-cut weight *w*^mc^ of the complexes. Square areas correspond to complexes and are superimposed in ascending order of *w*^mc^. At the left, the brain regions included in the complexes with high *w*^mc^ values (top 1, i.e., the main complex, to top 11) are enlarged. **c** The coreness values are plotted for each major brain region. The regions above the dashed line, which indicates the upper quartile of coreness values for all regions, correspond to the enlarged regions in **b**. **d** The inter-region connection matrix when bidirectionality is ignored, i.e. the mean of the original connection matrix *W* and its transpose *W^T^*, (*W* + *W^T^*)/2. **e** Structure of complexes. At the left, the brain regions included in the complexes with high *w*^mc^ values (top 1, i.e., the main complex, to top 14) are enlarged. **f** The coreness values are plotted for each major brain region. The regions above the dashed line, which indicates the upper quartile of coreness values for all regions, correspond to the enlarged regions in **e**.

#### Brain regions included in complexes

We extracted complexes in the mouse connectome using the proposed method. The extracted complexes are visualized in Fig. 7b. In Fig. 7b, the rows and columns of the connection matrix (Fig. 7a) are sorted according to the structure of the complexes in the same way as in Fig. 6. The color coding, which changes from blue to yellow, represents the value of the min-cut weight *w*^mc^ of the complexes. See Supplementary Table 1 for specific regions names and the detail values of *w*^mc^. On the left side of Fig. 7b, the brain regions included in the complexes with high *w*^mc^ values are enlarged. Specifically, the regions included in the complexes with the highest (a main complex) to the 11th highest *w*^mc^ are extracted. The 11th highest *w*^mc^ corresponds to the upper quartile of the coreness value for all regions.

In the following, we describe the regions that constitute the complexes with high *w*^mc^. In doing so, we do not distinguish between the left and right brains, because the extracted complexes were perfectly symmetric. That is, when a region on one side was included in a complex, the corresponding region on the opposite side was also included in the complex.

We observed that many regions in the cerebral cortex are included in top complexes (complexes with high *w*^mc^). In particular, mainly the isocortical regions constitute the first through third complexes. The only exceptions are the claustrum (CLA) and the basolateral amygdalar nucleus (BLA) in the cortical subplate, which are included in the third complex. The 4th to 9th complexes consist of the regions listed above plus other isocortical and thalamic regions, and the lateral parts of the entorhinal cortex (ENTl) in the hippocampal formation. The 10th and 11th complexes further includes some regions in the isocortex, olfactory areas, cortical subplate and pallidum. The regions in the other major regions are not included in the 1st to 11th complexes.

Thus, the regions included in the complexes with the highest *w*^mc^ are not evenly distributed among all major regions, but are rather concentrated in the cortical (particularly isocortical) regions and thalamic regions. We can confirm the unevenness among the major regions from the coreness values (Fig. 7d, Supplementary Table 2). Regions in the isocortex have particularly high coreness values (i.e., they are included in complexes with high *w*^mc^). Also, regions in the thalamus have high coreness values. Other regions with high coreness values are the CLA and BLA in the cortical subplate, followed by ENTl in the hippocampal formation, and some regions in the olfactory areas, cortical subplate and pallidum. On the other hand, regions in the other major regions have low coreness values. In particular, regions in the cerebellar cortex and cerebellar nuclei have much lower coreness values.

These results suggest that there appears to be a good correspondence between whether or not a region is included in complexes with high *w*^mc^ and whether or not a region is considered important for consciousness. For example, the isocortex and thalamus are considered essential to consciousness, whereas the cerebellar cortex and cerebellar nuclei do not directly contribute to consciousness ^1,19,33^. Other than the isocortical and thalamic regions, the CLA in the cortical subplate has long been associated with consciousness^47^. We discuss the relationship between consciousness and the regions included or not included in the top complexes in detail in Discussion.

#### Large difference in complexes when bidirectionality is ignored

Next, we investigated how the results change when the bidirectionality of the connections is ignored, i.e., the direction of connections is ignored and only the summed strength of connections is considered, as is in the example in Figs. 6d–f. Figure 7d shows the symmetrized connection matrix, based on which the complexes are extracted.

Let us first mention that similar to the case when considering bidirectionality, the results are symmetric between the left and right brains. That is, one of the following two conditions is satisfied: (1) as is in the case when considering bidirectionality, if a region on one side was included in a complex, the corresponding region on the opposite side was also included in the complex; or (2) if a region on one side was included in a complex *S*, the corresponding region on the opposite side was included in another complex with the same strength of bidirectional connections as that of *S*. We therefore do not distinguish between the left and right brains in the following.

Figure 7e represents the extracted complexes. See Supplementary Table 1 for specific regions names and the detail values of *w*^mc^. In the left side of Fig. 7e, the brain regions included in the complexes with the highest to 14th values of *w*^mc^ is enlarged. The 14th highest *w*^mc^ corresponds to the upper quartile of the coreness value for all regions. By comparing the brain regions in the top complexes shown in Fig. 7e with Fig. 7b, we can see that these are largely different in the sense that the brain regions in the top complexes are evenly distributed in almost all of the major brain regions when bidirectionality is ignored but are included in particular major regions such as the isocortex or thalamus when bidirectionality is considered. In fact, regions in all major regions except the cerebellar cortex are included in the complex with second highest *w*^mc^ when bidirectionality is ignored. We then investigate the change by ignoring bidirectionality using the coreness values (Fig. 7f, Supplementary Table 2). We observed that the difference among the major brain regions becomes small when bidirectionality is ignored. The maximum values of coreness are equal for many major regions. This reflects the fact that the regions included in the complex with large *w*^mc^ are evenly distributed in many major regions. Also, the median coreness values (represented by white circles in Fig. 7f) are equal for many major regions.

From Fig. 7f, we can see that there are two regions that have a particularly high coreness value, namely the frontal pole (FRP) in the isocortex and the caudoputamen (CP) in the striatum. The high coreness of these two regions is due to the strong connection from CP to the FRP. On the other hand, when bidirectionality is considered, the coreness value of CP is low because the connection in the opposite direction, FRP to CP, is weak.

To directly compare the two cases, namely when bidirectionality is considered or ignored, we made a scatter plot of coreness in Fig. 8, (1,2) or (2,1) panel. (See also the network diagram that compares the two cases in Supplementary Fig. 3). We can see that the distributions in the two cases are very different: regions with high coreness values when bidirectionality is considered do not necessarily have high coreness values when bidirectionality is ignored, and vice versa.

**Figure 8:**
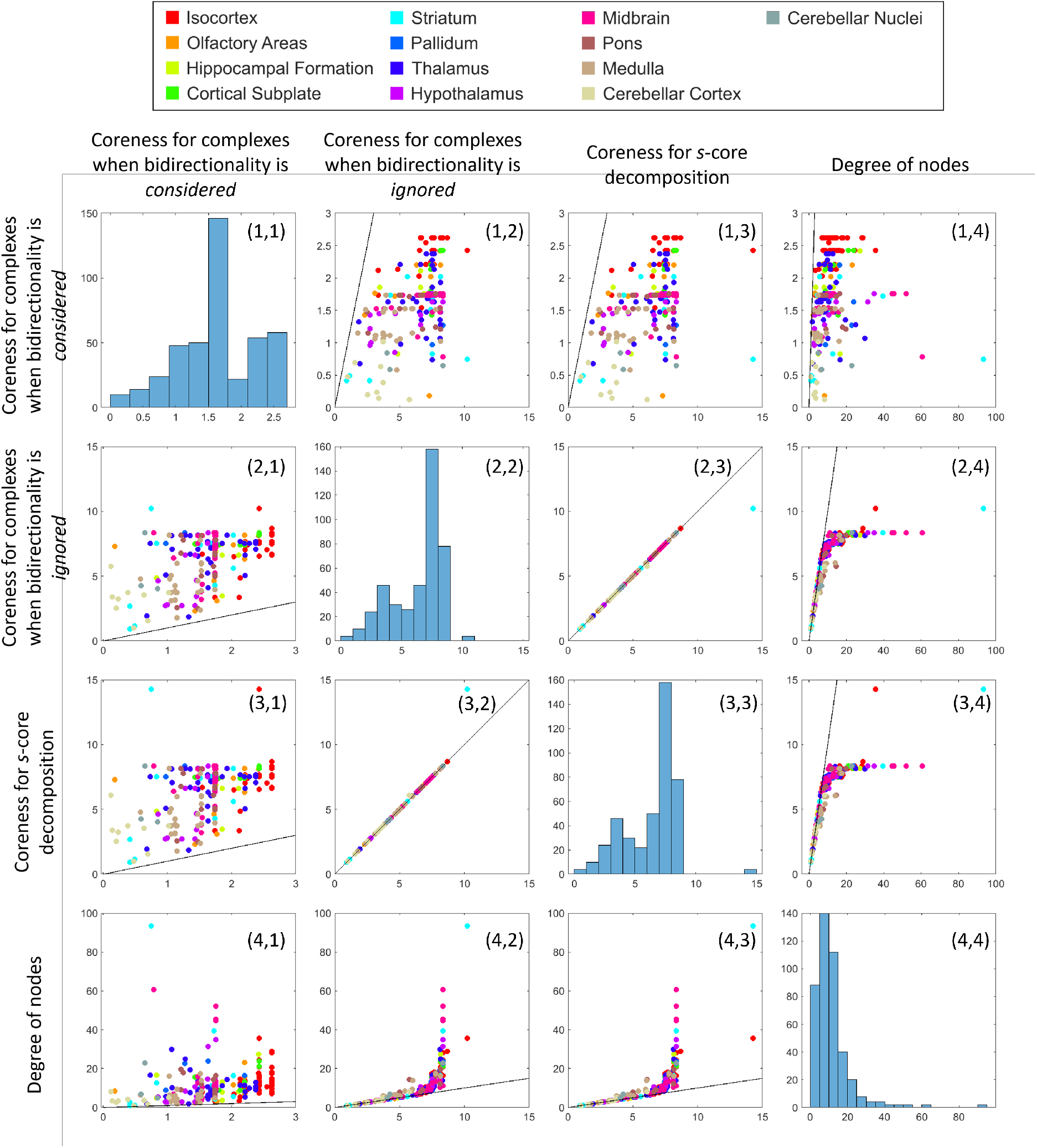
Histograms and scatter plots of coreness values and degree of nodes. Histograms of the coreness values and degree of nodes appear along the diagonal. Scatter plots of pairs of coreness value or degree of nodes appear in the off diagonal. Color of the points indicates major brain regions. The line in each scatter plot is the identity line (*y* = *x*).

Thus, if we ignore the bidirectionality of connections, the results change drastically; the complexes no longer necessarily consist of regions presumably essential to consciousness. This suggests that considering the bidirectionality of connections is important in associating the network core complexes with consciousness.

### Comparison with other existing methods

To further assess the significance of considering bidirectionality, we compare the proposed method with other existing methods that do not take account of bidirectionality. We first consider *s*-core decomposition ^43–46^, one of the most popular methods for extracting network cores. As we mentioned in the toy network analysis, *s*-core decomposition does not consider bidirectionality of connections and *s*-cores become identical to complexes when bidirectionality is ignored under a certain mathematical condition (See Supplementary Text 1 for details). In the mouse connectome case, this condition does not hold exactly, but almost does, and the obtained *s*-cores are almost the same as the complexes when bidirectionality is ignored. We can see that the coreness values for *s*-core decomposition (Coreness for *s*-core decomposition is defined in the same way as for complexes; see Methods) and those for the complexes when bidirectionality is ignored are almost identical (Fig. 8, (2,3) or (3,2) panel). Since for complexes the difference in coreness values among the major regions is small when bidirectionality is ignored (Fig. 7f), the difference is accordingly also small for s-cores. This means that the *s*-cores with a high *s* do not necessarily consist of regions in particular major regions, and therefore do not consist mainly of regions considered essential to consciousness.

Next, we investigated whether the complexes with strong bidirectional connections simply consist of the brain regions with high degree, i.e., network hubs^48,49^. The degree of a node is the sum of weights of edges connecting to it, irrespective of direction (Eq. (12) in Methods), and the network hubs are nodes with high degree. If the complexes with strong bidirectional connections consist of the hub regions, this means that bidirectionality does not matter to the extraction of complexes. We observed that the complexes when bidirectionality is considered do not necessarily consist of regions with high degree. In Fig. 8, (1,4) or (4,1) panel, we can see that the coreness values for the complexes when bidirectionality is considered and the degree are only weakly correlated: many brain regions with high coreness values have low degree. In contrast, the coreness for the complexes when bidirectionality is ignored (and the coreness for *s*-core decomposition) corresponds well to the degree, especially around the lower degree range (Fig. 8, (2,4) or (4,2) panel). Thus, these results indicate that the consideration of bidirectionality in the proposed method enabled us to extract core structures in the mouse connectome that cannot be extracted by simple degree-based methods.

## Discussion

In this study, we proposed a method to find the network cores, called “complexes”, that consist of strong bidirectional connections. If we search for complexes by brute force, computation time grows exponentially with the number of nodes. To solve this problem, we introduced a fast and exact algorithm proposed in our previous study, hierarchical partitioning for complex search (HPC)^41^. The HPC algorithm reduces the computation time to polynomial time and enables the analysis of large networks consisting of up to several thousand nodes in a practical amount of time. By utilizing HPC, we extracted complexes in a mesoscale, whole-brain mouse connectome consisting of 426 regions^42^, with the aim of identifying subnetworks in the brain relevant for consciousness. We found that complexes with strong bidirectional connections include many brain regions that have been considered essential for consciousness in previous studies. We also found that if bidirectionality is ignored, the brain regions included in the complexes with strong connections are evenly distributed in major brain regions regardless of whether or not they are relevant for consciousness. These results indicate that bidirectionality may be the key that characterizes the regions essential for consciousness.

### Correspondence between complexes and essential regions for consciousness

In this subsection, we discuss the relationship between complexes and the regions essential for consciousness. We first discuss in detail the brain regions with high coreness, i.e., the regions included in the complexes with strong bidirectional connections, and then the brain regions that are not included in such strong complexes.

First, many regions in the cerebral cortex, especially the isocortical regions, have high coreness. Previous studies suggested that bidirectional interaction among isocortical regions is essential for consciousness ^1,34,37^. In addition to the isocortical regions, the claustrum (CLA) in the cortical subplate also has high coreness. Francis Crick speculated that the CLA is the seat of consciousness, and that metaphorically speaking it plays the role of the conductor that orchestrates the brain^47^. In fact, recent studies in mice suggest that the CLA is involved in the control of arousal and sleep levels^50,51^. The CLA is also suggested to have a role in salience processing and attention control^52–57^, and might therefore be involved in selecting what comes to one’s conscious perception.

As for the subcortical regions, many thalamic regions also have high coreness. It is suggested that the thalamo-cortical loop - a circuit composed of the thalamus and cortical regions - is important for consciousness^58–61^.

As we discuss above, the brain regions with high coreness seem to correspond well with the regions that are considered essential to consciousness. However, the brain regions with high coreness also include some regions which have not yet been shown to be relevant to consciousness. A notable example is the basolateral amygdalar nucleus (BLA) in the cortical subplate. The BLA has the same coreness as the CLA, which is highest excluding the isocortical regions. The BLA is thought to be critical for emotion (positive and negative valence) and to mediate conditioning both for reward and fear^62^. To our knowledge, however, the relationship between the BLA and consciousness is little understood (e.g., whether the BLA directly contributes to subjective experience of emotions). Further investigation of such brain regions would be useful.

In addition to investigating whether the regions with high coreness are relevant to consciousness, it is also important to investigate the converse, namely whether the regions with low coreness, which are very weakly bidirectionally connected, are presumably irrelevant to consciousness. Notably, we found that all nodes in the cerebellar cortex and cerebellar nuclei have low coreness. It is well known that the cerebellum does not directly contribute to consciousness^3,4^ even though it has much more neurons than the cerebrum. We also found that the regions in the midbrain, medulla, and pons - the major regions which constitute the brainstem - have low coreness. Although the brainstem is important for enabling consciousness, it is not thought to contribute directly to conscious experience, in the same way that the heart is important for enabling consciousness but does not contribute directly to conscious experience. These are called background conditions ^1^.

Taking our results together, we have found that (1) brain regions presumably essential to consciousness have high coreness - that is, they are included in complexes with strong bidirectional connections; and that (2) brain regions presumably irrelevant to consciousness have low coreness, meaning that the regions are only weakly bidirectionally connected to other regions.

### Significance of considering bidirectionality

We here discuss how considering bidirectionality affects the results of the complexes and its importance in relating the complexes with the locus of consciousness. As we have seen in Results, when bidirectionality is ignored, the structure of the complexes largely differs from that when bidirectionality is considered. One large difference is that the difference in coreness between major regions is smaller when bidirectionality is ignored. Regions in the cerebellum (cerebellar cortex and cerebellar nuclei) and brainstem (midbrain, pons and medulla), which have smaller coreness than other regions such as the isocortex and thalamus when bidirectionality is considered, have similarly high coreness to these regions when bidirectionality is ignored. As mentioned in the previous section, these regions have not been so far considered to directly contribute to consciousness ^1,3,4^. Another large difference is that the caudoputamen (CP) in the striatum, which is not included among complexes with large *w*^mc^ when bidirectionality is considered, forms the main complex when bidirectionality is ignored. The striatum, more broadly the basal ganglia, is not thought to contribute directly to consciousness^5,39^ (but see^63,64^).

Thus, when bidirectionality is ignored, regions both relevant and irrelevant to consciousness are evenly included in the strong complexes. Thus, the seemingly good correspondence between complexes and regions relevant to consciousness we identified when considering bidirectionality is largely lost.

### Comparison with other core extraction methods in terms of bidirectionality

In the literature, a variety of methods have been applied to connectomes to extract network cores in which elements are densely connected to each other. In what follows, in terms of bidirectionality of connections, we compare complexes with three representative methods for core extraction, namely *s*-core decomposition, network hubs, and modularity maximization.

In this study, we first compared *s*-core decomposition with the complexes. *s*-core decomposition is a representative method which has been widely applied to connectomes of various species^43,44,46^, and the extracted cores are related to certain functions. We showed that the *s*-cores extracted from the mouse connectome largely differ to the complexes when bidirectionality is considered, but are almost identical to the complexes when bidirectionality is ignored. This means that the consideration of bidirectionality enabled us to reveal core structures that cannot be revealed by *s*-core decomposition.

We next compared network hubs^48,49^ with the complexes. Previous studies showed that the brain network contains cores in which hubs (high-degree nodes) are densely connected to each other (called “rich-clubs”)^45,46,49^. We showed that in the mouse connectome the complexes with strong bidirectional connections included not only high-degree nodes but many low-degree nodes (Fig. 8). This means that the core structures revealed by the complexes largely differ to the structures that can be revealed by hub-based methods.

Finally, we discuss modularity maximization, which is also widely used in connectome analysis^65^. Similar to the proposed method, modularity maximization is a method used to extract subnetworks with dense connections. Its objective is, however, qualitatively different from that of the proposed method. The objective of modularity maximization is to partition a network into non-overlapping cores (called modules or communities) with dense internal connections, and not to decompose a network hierarchically as for complexes. This difference in objectives hampers direct quantitative comparison of the two methods by experiments. We therefore confined ourselves here to a qualitative comparison in terms of bidirectionality. The mathematical formulation of modularity maximization is suitable for undirected networks^65^. It is therefore impossible to consider the direction of connections and hence bidirectionality. However, a variant of the modularity maximization methods considers the direction of connections when defining the density of connections^66^. This variant does not consider bidirectionality, however, and extracted modules do not therefore necessarily consist of bidirectional connections, i.e., modules have fully feed-forward structures.

As exemplified above, the core extraction methods in wide current use for connectome analysis do not consider the bidirectionality of connections. Thus, we conclude that the main result of the present paper, which has revealed the correspondence between the network cores of the brain and consciousness, can only be achieved by methods such as the proposed method which take account of the bidirectionality of connections.

### Number of main complexes

In general, since main complexes are “local” maxima in terms of *w*^mc^, there can be multiple main complexes in a network as we mentioned in Results. An extreme example occurs when a network consists of two mutually disconnected modules: in this case, there will be two (or more) main complexes. The presence of multiple main complexes in a network indicates that the network consists of multiple weakly coupled modules.

In the mouse connectome, there are five main complexes when bidirectionality is considered, albeit that we only mentioned one of them in Results. The main complex we mentioned has the highest *w*^mc^ of all main complexes (and of all subnetworks by definition) and is largest in size among all main complexes. The other main complexes, which we did not mention, have low *w*^mc^ and consist of only two regions, i.e., are minimum in size. This means that the mouse connectome can be almost considered to consist of one large module.

Thus, we can ascertain modular structure using complexes. In contrast, when we use *s*-core decomposition, this cannot be ascertained. Consider a network consisting of two densely connected parts, as shown in Supplementary Fig. 4. In this case, s-core decomposition extracts the entire network as a single core and does not extract the modular structure in this network. The proposed method, on the other hand, extracts two modules as two main complexes. This is because *s*-core decomposition uses only local information, i.e., degree of nodes, and cannot consider the global structure of a network as a whole. On the other hand, the proposed method uses the global information of networks, min-cuts, which allows us to extract the modular structure.

### Limitations of this study and future direction

We searched for complexes using a method proposed in our previous study, hierarchical partitioning for complex search (HPC)^41^. The computation time of HPC increases only polynomially with the number of the nodes, which is much smaller than the exponential increase in brute force search. This enables us to find complexes in a network with several thousand nodes in a practical amount of time. However, to find complexes in networks with more nodes (*N* ≫ 2,000–3,000), a further speeding-up is required. One possible solution is the use of approximation algorithms for min-cut search^67^ instead of the exact algorithm, the Hao-Orlin algorithm^68^, which we used in this study.

In this study, we discussed how complexes extracted from the mouse connectome, which consists of anatomical connections, are related to consciousness. We should note, however, that it is not the anatomical connections themselves that are directly responsible for conscious experiences at a particular time, but rather interactions between brain regions that result from the brain activity^9–17^. The location of bidirectional interactions changes from time to time, and the brain regions that mediate consciousness can also change accordingly^1,33^. To capture such dynamic change in consciousness, future research should therefore aim to extract complexes from functional or causal networks constructed by quantifying interactions using brain activity data. The relationship between anatomical and functional networks is not as simple as a one-to-one correspondence. It is empirically known, however, that there are some similarities between them^69–71^, as would be naturally expected from the fact that anatomical connections are the basis for interactions between brain regions. We therefore expect that complexes extracted from functional networks could be similar to the complexes extracted from the anatomical network.

## Methods

### Strength of bidirectional connections

Here, we propose a way of quantifying how strongly two parts of a graph are bidirectionally connected. We consider a directed graph *G*(*V,E*), where *V* and *E* are the node set and the edge set, respectively. For a bi-partition of the node set *V*, (*V*_L_, *V*_R_), there are two types of edges that connect *V*_L_ and *V*_R_ depending on its direction. One is the set of edges outgoing from *V*_L_ to *V*_R_:

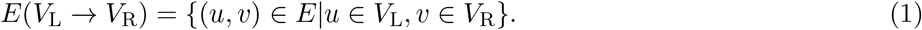

The other is the set of the edges incoming to *V*_L_ from *V*_R_ (or equivalently, outgoing from *V*_R_ to *V*_L_):

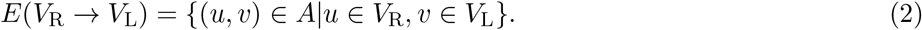

When we ignore the directions of the connections between *V*_L_ and *V*_R_, the simplest way of quantifying the strength of the connections is to add up all the weights of the edges that connect *V*_L_ and *V*_R_ regardless of their directions:

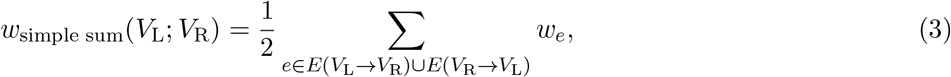

where *w_e_* represent the weight of the edge *e*. The factor 2 in the denominator is for consistency with the strength of bidirectional connections, as explained later.

On the other hand, when we consider the bidirectionality of connections between *V*_L_ and *V*_R_, we first separately add up the weight of the edges for each direction,

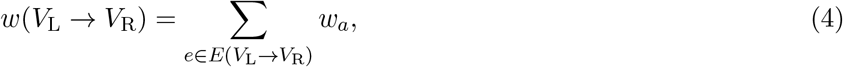

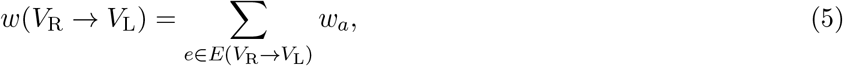

and then define the strength of bidirectional connections as their minimum:

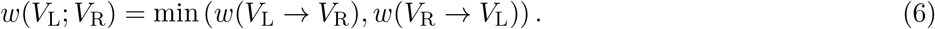

With this definition, if two parts of a network are only connected unidirectionally, as in Fig. 2a, the strength of bidirectional connections *w*(*V*_L_; *V*_R_) is 0, which means that the two parts are considered to be “disconnected” bidirectionally. In Fig. 2b, the connection from one part to the other part is strong (3) but that in the other direction is weak (1). Consequently, the strength of bidirectional connections is low (*w*(*V*_L_; *V*_R_) = 1). In Fig. 2c, the connections in both directions are strong (2) and the strength of bidirectional connections is high (*w*(*V*_L_; *V*_R_) = 2). If we ignore the directionality of connections and add up the edge weights in the two directions (*w*_simple sum_), the strength of connections is evaluated as 2 in all three cases.

The two measures *w*(*V*_L_; *V*_R_) and *w*_simple sum_, with and without considering bidirectionality, are equal to each other when connections are symmetric (*w*_(*u,v*)_ = *w*(_*v,u*_)).

### Minimum cut

#### Definition of minimum cut

A cut of a graph *G*(*V*, *E*) is called a minimum cut if the strength of the connections for the cut is not higher than that of any other cut. More formally, a minimum cut 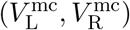 is defined as follows:

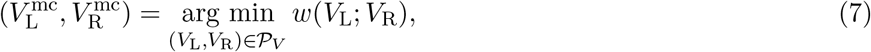

where 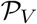 denotes the set of all bi-partitions of *V*. We denote the weight of the minimum cut 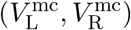 of a graph *G* as

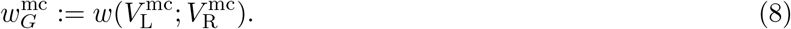

#### Fast and exact algorithms for searching for min-cuts

We defined a measure of strength of bidirectional connections as in Eq. (6). Although this definition is different from the canonical definition of a graph cut weight for directed graphs, the minimum cut problems for the two definitions are equivalent (Supplementary Text 2). Therefore, we can use a well-established algorithm to solve the minimum cut problem. In this study, we utilize the Hao-Orlin algorithm^68^. Its time complexity is *O*(|*V*||*E*|log(|*V*|^2^/|*E*|), where |*V*| and |*E*| are the number of nodes and edges, respectively.

### Complex

In this section, we introduce the definition of a complex^40,41,72^. We also introduce a main complex, which is a stronger definition of a complex^40,41,72^.

To formally define complexes, we need to introduce the concept of an induced subgraph. Let *G* be a graph consisting of node set *V* and edge set *E*, and let *S* ⊆ *V* be a subset of nodes. Then, an induced subgraph *G*[*S*] is the graph consisting of all the nodes in *S* and all the edges connecting the nodes in *S*. The min-cut weight of *G*[*S*] is denoted by 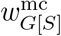. We are now ready to define complexes.

#### Definition 1 (Complex)

*An induced subgraph G*[*S*] (*S* ⊆ *V*) *is called a complex if it satisfies* 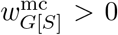 *and* 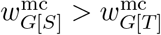 *for any subset T that is a superset of S* (*T* ⊃ *S and T* ⊆ *V*).

A schematic explanation of the definition of a complex is shown in Fig. 4. In this schematic, we consider induced subgraphs of a graph *G* consisting of ten nodes {*A, B*,…, *J*}. An induced subgraph *G*[{*E, F, I, J*}] is a complex because it has greater *w*^mc^ than any induced subgraph of *G* that is its supergraph (e.g., *G*[{*B, E, F, I, J*}] and *G*[{*D,E,F,H,I, J*}]).

The whole graph *G* is a complex if it satisfies 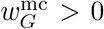 by definition. We define *w*^mc^ = 0 for single nodes because we cannot consider partitions of a single node. Therefore, single nodes cannot be complexes.

An induced subgraph is called a main complex if its min-cut weight *w*^mc^ is larger than those of any induced subgraphs that are its supergraphs, and is also larger than or equal to those of any induced subgraphs that are its subgraphs. That is, a complex is called a main complex if its min-cut weight *w*^mc^ is larger than or equal to those of any induced subgraphs that are its subgraphs.

#### Definition 2 (Main complex)

*A complex is called a main complex if it satisfies* 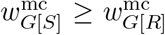 *for any subset R of S* (*R* ⊂ *S*).

A schematic explanation of the definition of main complexes is shown in Fig. 4. An induced subgraph *G*[{*E, F, I, J*}] is a main complex because it is a complex and has greater *w*^mc^ than any induced subgraph that is its subgraph (e.g., *G*[{*F, J*}] and *G*[{*E, F, I*}]).

### Hierarchical Partitioning for Complex Search

If we search for complexes by brute force, computation time increases exponentially with the number of nodes. Therefore, when the number of nodes in the network exceeds several tens, it becomes practically impossible to identify the complexes. On the other hand, using the algorithm Hierarchical Partitioning for Complex Search (HPC), which we proposed in a previous study^41^, the computation time increases only polynomially with the number of nodes. HPC is an exact method that does not use approximations and can extract all complexes without any omissions or misidentifications. This makes it possible to extract all complexes from a network consisting of several thousand nodes in a practical computation time. An actual computation time evaluated by a simulation is shown in Supplementary Fig. 1.

In what follows in this subsection we write the induced subgraph *G*[*S*] for a node subset *S* as *S* for simplicity of notation.

HPC primarily consists of two steps. The first is listing candidates of (main) complexes. HPC narrows down candidates for (main) complexes by hierarchically partitioning a network. The second step is screening the candidates to find (main) complexes.

In the first step, HPC hierarchically partitions a network with min-cuts (Fig. 5). HPC starts by dividing the whole network with its min-cut, and then repeatedly divides the subnetworks with their min-cuts until the entire network is completely decomposed into single nodes. This procedure in HPC is summarized as follows:

1. Find the min-cut (*V*_L_, *V*_R_) of the whole network *V* and divide the whole network *V* into the two subnetworks *V*_L_ and *V*_R_.
2. Find the min-cuts of the subnetworks found in the previous step, *V*_L_ and *V*_R_, and divide them into (*V*_LL_, *V*_LR_) and (*V*_RL_, *V*_RR_), respectively.
3. Repeat this division until the whole network is decomposed into single nodes.

After the procedure above, we obtain the set of hierarchically partitioned subnetworks, that is, *V*, *V*_L_, *V*_R_, *V*_LL_, *V*_LR_, *V*_RL_, *V*_RR_, and so on. We consider all the set of subnetworks

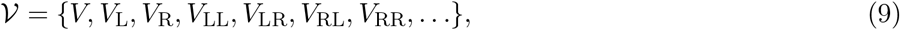

excluding single nodes. Then, the following theorem holds.

#### Theorem 3.

*Any complex S* ⊆ *V belongs to* 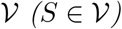.

Thus from this theorem, 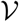 can be seen as the set of candidates of complexes. The theorem is based on satisfaction of a mathematical property “monotonicity” by the strength of bidirectional connections (Eq. (6)). Let us consider the strength of bidirectional connections *w*(*S*; *T*) between two subsets of nodes *S* and *T*. If we then add another set of nodes *U* to *S*, the strength of bidirectional connections does not decrease. That is,

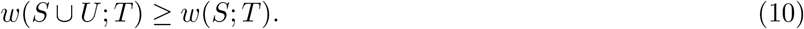

Also, if we add *U* to *T*, *w*(*S*;*T* ∪ *U*) ≥ *w*(*S*;*T*). This inequality means that the strength of bidirectional connections monotonically increases as nodes are added. We call this property “monotonicity”. By using monotonicity, we can easily show that a subnetwork cannot be a complex if it straddles the boundary of a min-cut of a subnetwork that contains it, and can prove Theorem 3 (see our previous work^41^ for the proof).

After the hierarchical partitioning procedure described above, in the second step, we need to check whether each candidate of complexes belonging to 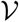 is actually a (main) complex or not in accordance with Def. 1. We can efficiently check this by taking advantage of the hierarchical (tree) structure. For more detail please see our previous work ^41^.

In general, a network can have multiple min-cuts. If this is the case, depending on which min-cut is used to divide a network in the hierarchical partitioning process, the candidate set of complexes 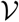 can vary. However, even though 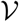 varies, the resulting complexes (and also main complexes) do not vary. This is because any of the candidate sets contains all (main) complexes independent of which min-cut is used. Therefore we do not have to care which of multiple min-cuts we select.

### Coreness of each node

Using the complexes and their *w*^mc^, we define a “coreness” of each node. When a node is included in complexes with high *w*^mc^, the coreness of that node is high, and conversely, when a node is included only in complexes with low *w*^mc^, the coreness of that node is low. Specifically, we define the coreness of a node *v* as *k_v_* if the node *v* is included in a complex with *w*^mc^ = *k_v_* but not included in any complex with *w*^mc^ > *k_v_*. Equivalently, we can define the coreness of a node *v* as the largest of the *w*^mc^ of all complexes containing the node *v*:

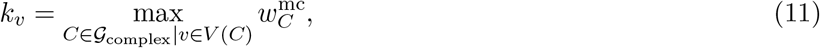

where 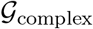 denotes the set of all complexes in the graph *G* and *V*(*C*) denotes the set of all nodes in the complex *C*.

In the same way, we can define a coreness for *s*-core decomposition: we define the coreness of a node *v* as *s* if node *v* is included in *s*-core but is not included in any *s*′-core with *s*′ > *s*.

### Degree of a node

We define the degree of a node *v* as the sum of the weights of all edges connecting *v* and other nodes, irrespective of the direction of edges:

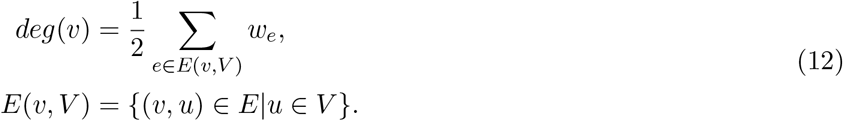

The factor 2 in the denominator is for consistency with the strength of connections: when we measure the strength of connections (Eq. (3)) between a node and the nodes connecting to it, it becomes equal to the degree of the node. This degree can be also regarded as the mean of in-degree and out-degree.

### Sorting rows and columns of a connection matrix according to the structures of complexes

In Figs. 6 b and 6h and Figs. 7 d and 7h, we sorted rows and columns of a connection matrix according to the hierarchical structures of the complexes. Here we explain this sorting process in detail.

To start, we sort the rows and columns in the order of the leaf nodes of the hierarchical structure obtained by hierarchical partitioning (Fig. 5). In the case of Fig. 5, the rows (columns) are sorted in the order of *A, B, E, F, C, D*, and *G*. We now explain in detail. At each step of the hierarchical partitioning process, we sort the nodes according to which of the two subnetworks (e.g., *V*_L_ or *V*_R_) they are classified in. Therefore, at the end of the process, nodes classified into the same groups until a late stage of the process are placed close to each other, whereas those classified into different groups at an early stage are placed away from each other. Since the hierarchical structure obtained by the hierarchical partitioning is the basis of the hierarchical structure formed by the complexes, the result is that nodes in the same complex with high *w*^mc^ are placed close to each other.

This is the rough flow of how the order of nodes is determined. This alone, however, is not enough to uniquely determine the order. There is still arbitrariness with regard to which of the two subnetworks (e.g., *V*_L_ or *V*_R_) comes first at each step of the process. To eliminate this arbitrariness, we chose to place the upstream subnetwork first. That is, for example, when the strength of the connections from *V*_L_ to *V*_R_ (Eq. (4)) is higher than that in the opposite direction (Eq. (5)), i.e., when *V*_L_ is located relatively upstream to *V*_R_, *V*_L_ is placed ahead of *V*_R_. If the strengths of the connections in the two directions are equal to each other, we arranged the rows (columns) so that their original order is maintained.

## Acknowledgments

This work was partially supported by JST ACT-X Grant Number JPMJAX20A6, JST CREST Grant Numbers JPMJCR1864 and JPMJCR15E2 including AIP challenge program, and JSPS KAKENHI Grant Numbers 18H02713 and 20H05712, Japan.

## Supplementary information

**Supplementary Text 1.** *s*-**core decomposition and its relation to the proposed method.**

**Supplementary Text 2. Equivalence between the min-cut considering bidirectionality and the canonical min-cut for directed graphs.**

**Supplementary Figure 1. Computation time of Hierarchical Partitioning for Complex Search.** Computation time was evaluated by a simulation. In the simulation, networks with different numbers of nodes were randomly generated. The weight of each edge was sampled from a uniform distribution in the interval (0, 1). The red circles and the red solid lines indicate the computation time of Hierarchical Partitioning for Complex Search and a fitted linear function (log_10_*T* = 3.419 log_10_ *N* — 7.463 (*T* ∝ *N*^3.419^)). The black triangles and black dashed lines indicate the computation time of the exhaustive search and a fitted exponential function (log_10_*T* = 0.5421*N* – 5.179). The simulation was done on a machine with an Intel Xeon Gold 5220 processor at 2.20 GHz. All the calculations were implemented in MATLAB 2019a.

**Supplementary Figure 2. A network with two main complexes.** There are two main complexes ({*D, E, F*} and {*G, H, I*}) and the complexes form a nested hierarchical structure with the two main complexes as peaks.

**Supplementary Figure 3. Network diagrams.** To directly compare the network structure revealed by the complexes when bidirectionality is considered and ignored, we plotted a network diagram. Each node represents a brain region. The color of each node indicates the major brain region in which the node is included. The size of each node indicates its degree. Each edge represents the connection between a node pair. The width of each edge is proportional to the edge weight (for visibility, only edges with the top 20% weight are shown). The color of each edge indicates the min-cut weight *w*^mc^. More specifically, if two nodes are included in the same complex with *w*^mc^ = *w*, then the edge between them is colored with the color corresponding to w. The color is overwritten in the ascending order of *w*^mc^. Therefore, if two nodes are included in a complex with high *w*^mc^, the edge color becomes yellowish, whereas if they are included only in complexes with low *w*^mc^, the edge color becomes bluish. We arranged the points so that the x-coordinate of each point is the same as the order of the rows (columns) in the sorted connection matrix in Fig. 7b, where bidirectionality is considered. More specifically, the brain region that is at the *i*-th row (column) of the sorted connection matrix in Fig. 7b is placed at the position of *x* = *i*. As a comparison, we set the y-coordinates of the points to be the same as the order of the rows (columns) in the sorted connection matrix in Fig. 7e, where bidirectionality is ignored. As described in Methods, we sorted the rows and columns of the connection matrix according to the structure of complexes, and also sorted them so that relatively upstream nodes come first and relatively downstream nodes come later. Therefore, points with a smaller or larger x-coordinate respectively correspond to relatively upstream or downstream regions. In (a), we draw only rightward edges, which start from the smaller x-coordinate and end at the larger x-coordinate. On the contrary, in (b), we draw only leftward edges, which start from the larger x-coordinate and end at the smaller x-coordinate. We can see that the nodes in the complexes with high *w*^mc^ are connected to each other by both rightward and leftward edges, which are indicated by yellowish color. In other words, the nodes are bidirectionally connected to each other. These nodes are mainly in the major regions such as the isocortex, hippocampal formation, cortical subplate and thalamus. The isocortical regions occupy the center of the complexes, i.e., the main complex. The thalamic regions are also located close to the center, but relatively upstream compared to the isocortical regions. On the other hand, the nodes at the left side of the figure have many edges going out but few edges coming in. In other words, these nodes are located upstream in the network. Conversely, the nodes at the right side of the figure have many edges coming in but few edges going out, i.e., these nodes are located downstream in the network.

**Supplementary Figure 4. Comparison of the proposed method with *s*-core decomposition. a** A network consisting of two densely connected parts. **b** If we apply *s*-core decomposition to the network in **a**, the entire network is extracted as the *s*_max_-core and the modular structure in this network is not revealed. **c** If we apply the proposed method, the two modules are extracted as two main complexes regardless of whether bidirectionality is considered or not.

**Supplementary Table 1. Sorted connection matrix, with the region names and the values of** *w*^mc^. The first sheet is the sorted connection matrix when bidirectionality is considered, which is the same as that in Fig. 7b. The second sheet is the sorted connection matrix when bidirectionality is ignored, which is the same as that in Fig. 7e.

**Supplementary Table 2. Coreness values of all regions.** In the first and second sheets, the coreness values when bidirectionality is considered and ignored, respectively, are listed in descending order.

