## Supplementary Text 1 for "Bidirectionally connected cores in a mouse connectome: Towards extracting the brain subnetworks essential for consciousness"

## Supplementary Text 1   $s$-core decomposition and its relation to the proposed method

Here we theoretically compare complexes and $s$-cores. To do so, let us first introduce the definition of $s$-cores and the algorithm for finding them.

#### Definition of $s$-cores

$s$-core decomposition[44,45] is a method to identify densely connected parts in a graph. It is a generalization of $k$-core decomposition[43–46]. $s$-cores are defined for "weighted" graphs, while $k$-cores are defined for "unweighted" graphs. Let us consider a weighted undirected graph $G$. An $s$-core is then defined as follows.

**Definition 4** ($s$-core)**.** *An $s$-core is a maximal subgraph of $G$ in which all nodes have a degree of at least $s$.*

Here "maximal" means that a subgraph is the largest one: we cannot add another node to the subgraph without violating the condition "all nodes have a degree of at least $s$".

We can hierarchically decompose a network into $s$-cores with different $s$ in a similar way as for complexes. A core includes another core with a larger $s$, and the inner core includes another core with an even larger $s$. We refer to the largest $s$ as $s_{\max}$ and the corresponding core as $s_{\max}$-core.

#### Algorithm for finding $s$-cores

We can find $s$-cores in a very simple way: recursively pruning the nodes with the minimum degree. The entire process is described as follows:

1. Find the node with the minimum degree in the entire graph $G$ and refer to the minimum degree as $s_{\text{temp}}$. The entire graph is an $s_{\text{temp}}$-core.
2. Recursively prune the nodes with the minimum degree while the degree is less than or equal to $s_{\text{temp}}$. If there are no more nodes to prune, then the remaining subgraph (if it is not empty) is a $s'$-core, where $s'$ denotes the minimum of the degrees of the nodes in the remaining subgraph. Note that $s' > s_{\text{temp}}$.
3. Set $s_{\text{temp}} = s'$.
4. Repeat Steps 2 and 3 until the graph becomes empty and we obtain all $s$-cores.

### Difference between complexes and $s$-cores

A simple difference is whether we can take into account the bidirectionality of the connections. When analyzing a directed graph with $s$-core decomposition, we need to ignore the direction of edges and treat the graph as an undirected one, or consider only either in-coming or out-going edges. We are therefore unable to take into account the bidirectionality of the connections.

Apart from this difference in directionality, we give a typical example where the results largely differ between the $s$-core decomposition and the proposed method. Consider a graph consisting of two densely connected parts, as shown in Supplementary Fig. 4. In this case, $s$-core decomposition extracts the entire graph as the $s_{\max}$-core and does not extract the modular structure in this graph. The proposed method, on the other hand, extracts two modules as two complexes. This is because $s$-core decomposition uses only local information, i.e., degree of nodes, and cannot consider a global structure of a graph as a whole. On the other hand, the proposed method uses the global information of graphs, min-cuts, which allows us to extract the modular structure.

#### A sufficient condition under which complexes and $s$-cores become the same

So far we have discussed the difference between the proposed method and $s$-core decomposition. Next, we discuss cases where the results of the proposed method and $s$-core decomposition become the same. We prove that complexes when bidirectionality is ignored and $s$-cores become identical under a certain mathematical condition: when all min-cuts used in the proposed method separate a single node from the others. In this case, the computational process of the proposed method and that of $s$-core decomposition are the same, and therefore we get the same results.

**Theorem 5.** *Consider complexes and $s$-cores of an undirected graph. If all min-cuts separate one node from the others in the hierarchical partitioning process (Fig. 5), complexes and $s$-cores become identical: a complex with $w^{\mathrm{mc}} = w$ is a $w$-core and an $s$-core is a complex with $w^{\mathrm{mc}} = s$. In particular, the main-complex corresponds to the $s_{max}$-core.*

*Proof.* When a min-cut in a graph separates one node from the others, the separated node is the node with the minimum degree (and the degree is the min-cut weight $w^{\mathrm{mc}}$). Therefore, when all min-cuts separate one node from the others in the hierarchical partitioning (HP) process, the sequence of hierarchical partitioning becomes the same as the sequence of recursive pruning in $s$-core decomposition. The condition for a candidate complex $S$ ($S \in \mathcal{V}$) to be a complex is that $w^{\mathrm{mc}}$ of $S$ is larger than those of the ancestor candidates in $\mathcal{V}$[41]. This condition is equivalent to the condition in which a subgraph appearing in the pruning process is an $s$-core for an $s$ ($s' > s_{\mathrm{temp}}$). Thus, complexes and $s$-cores are identical. In particular, the main complex corresponds to the $s_{\mathrm{max}}$-core because the main complex is a complex with the largest $w^{\mathrm{mc}}$. □

In the toy network in Results, the condition holds and the complexes when bidirectionality is ignored and the $s$-cores are therefore identical. In the mouse connectome, the condition largely but not completely holds (i.e., most but not all of the min-cuts appearing in the hierarchical partitioning process separate a single node from the others), and the results of the proposed method when bidirectionality is ignored and $s$-core decomposition are almost the same (Fig. 8).

A min-cut tends separate a single node from the others when a graph is densely connected and has a low variability of weights (i.e., a graph has a nearly uniform structure). Thus, complexes and $s$-cores in such cases tend to be identical.
