## Supplementary Text 2 for "Bidirectionally connected cores in a mouse connectome: Towards extracting the brain subnetworks essential for consciousness"

### Supplementary Text 2 Equivalence between the min-cut considering bidirectionality and the canonical min-cut for directed graphs

We show that the min-cut defined by Eq. (7) in the main text is equivalent to the canonical definition[68] of min-cut for directed graphs. The min-cut defined by Eq. (7) in the main text can be transformed as follows.

$$
\begin{aligned}
&\min_{(V_\mathrm{L}, V_\mathrm{R}) \in \mathcal{P}_V} w(V_\mathrm{L}; V_\mathrm{R}), \\
=&\min_{(V_\mathrm{L}, V_\mathrm{R}) \in \mathcal{P}_V} \left\{ \min\left( w(V_\mathrm{L} \to V_\mathrm{R}), w(V_\mathrm{R} \to V_\mathrm{L}) \right) \right\}, \\
=&\min_{(V_\mathrm{L}, V_\mathrm{R}) \in \mathcal{P}_V} w(V_\mathrm{L} \to V_\mathrm{R}), \\
=&\min_{V_\mathrm{L} \subseteq V, V_\mathrm{L} \neq \emptyset} w(V_\mathrm{L} \to V \backslash V_\mathrm{L}),
\end{aligned}
$$

which is equivalent to the canonical min-cut for directed graphs.
