## Supplementary figures and images for "Bidirectionally connected cores in a mouse connectome: Towards extracting the brain subnetworks essential for consciousness"

### Supplementary Figure 1

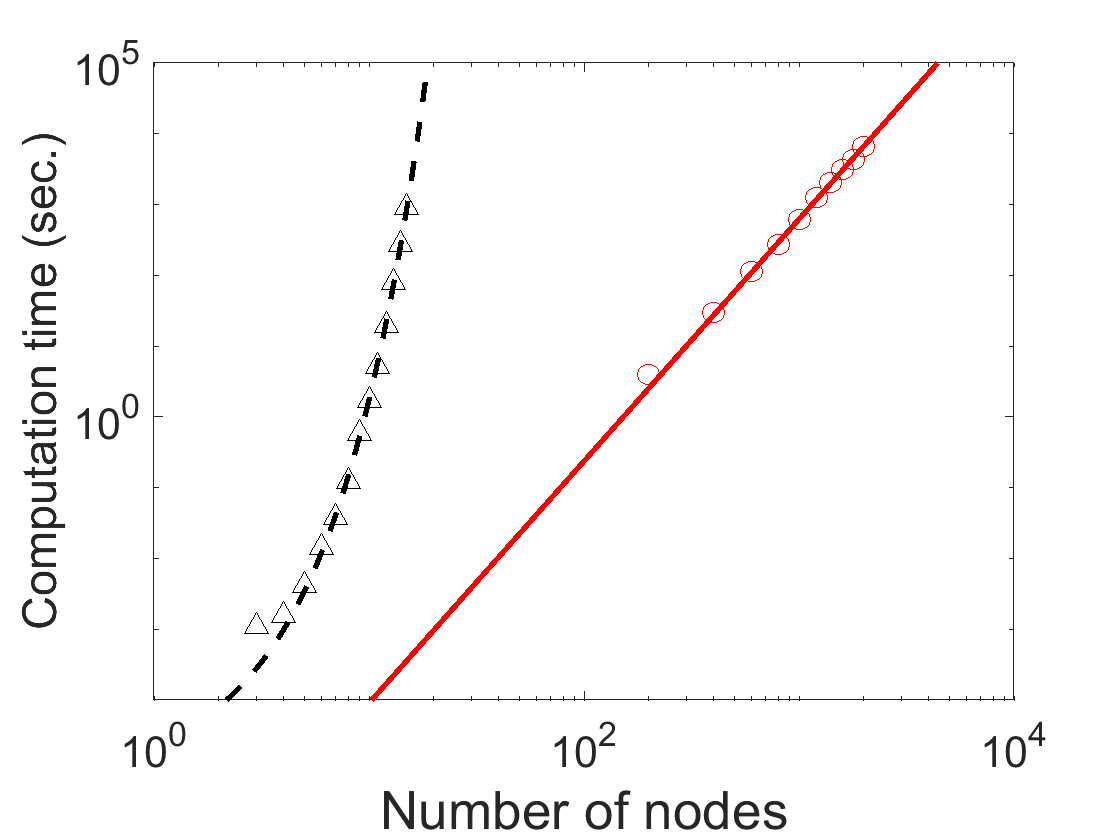

### Supplementary Figure 2

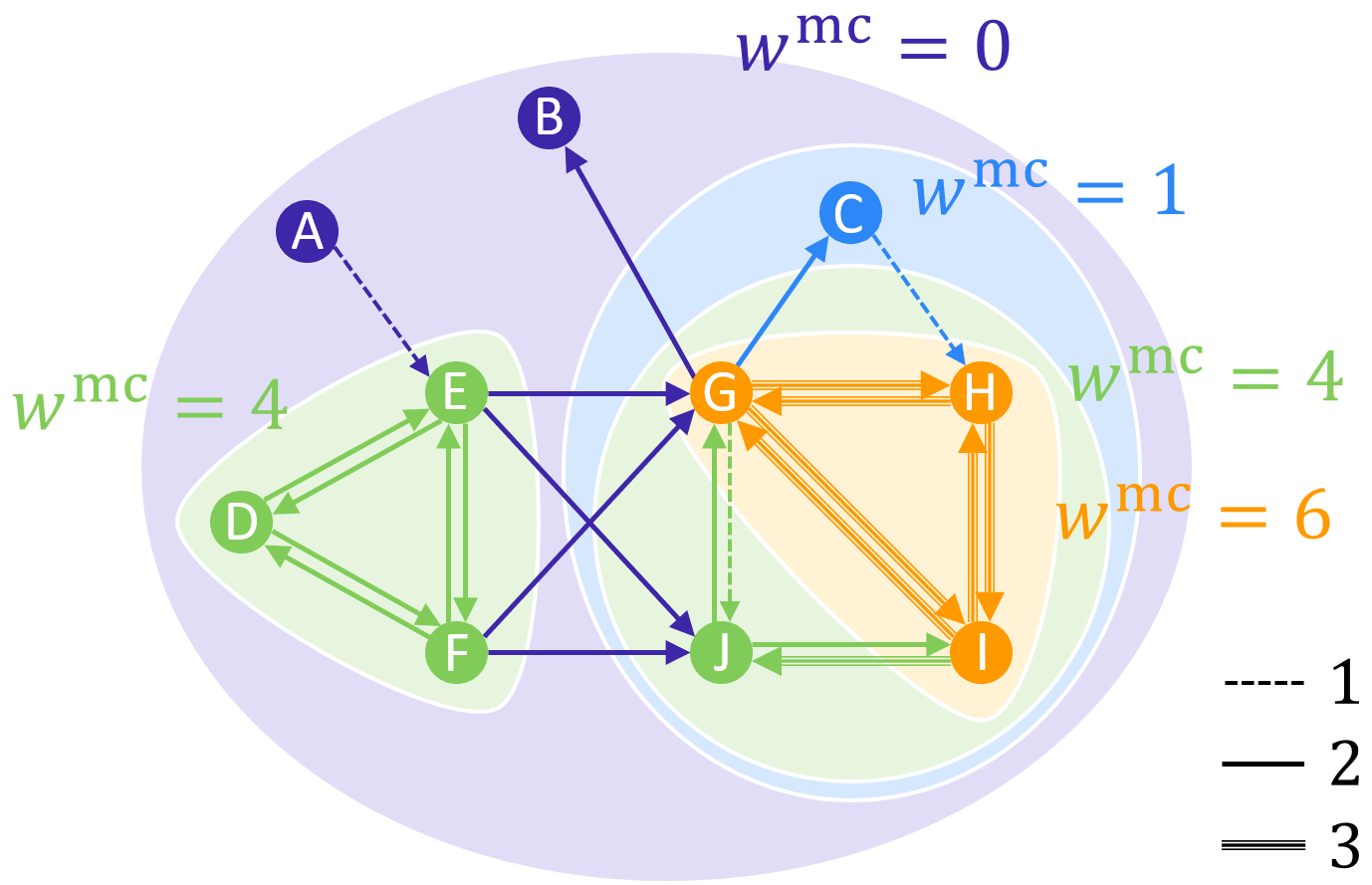

### Supplementary Figure 3

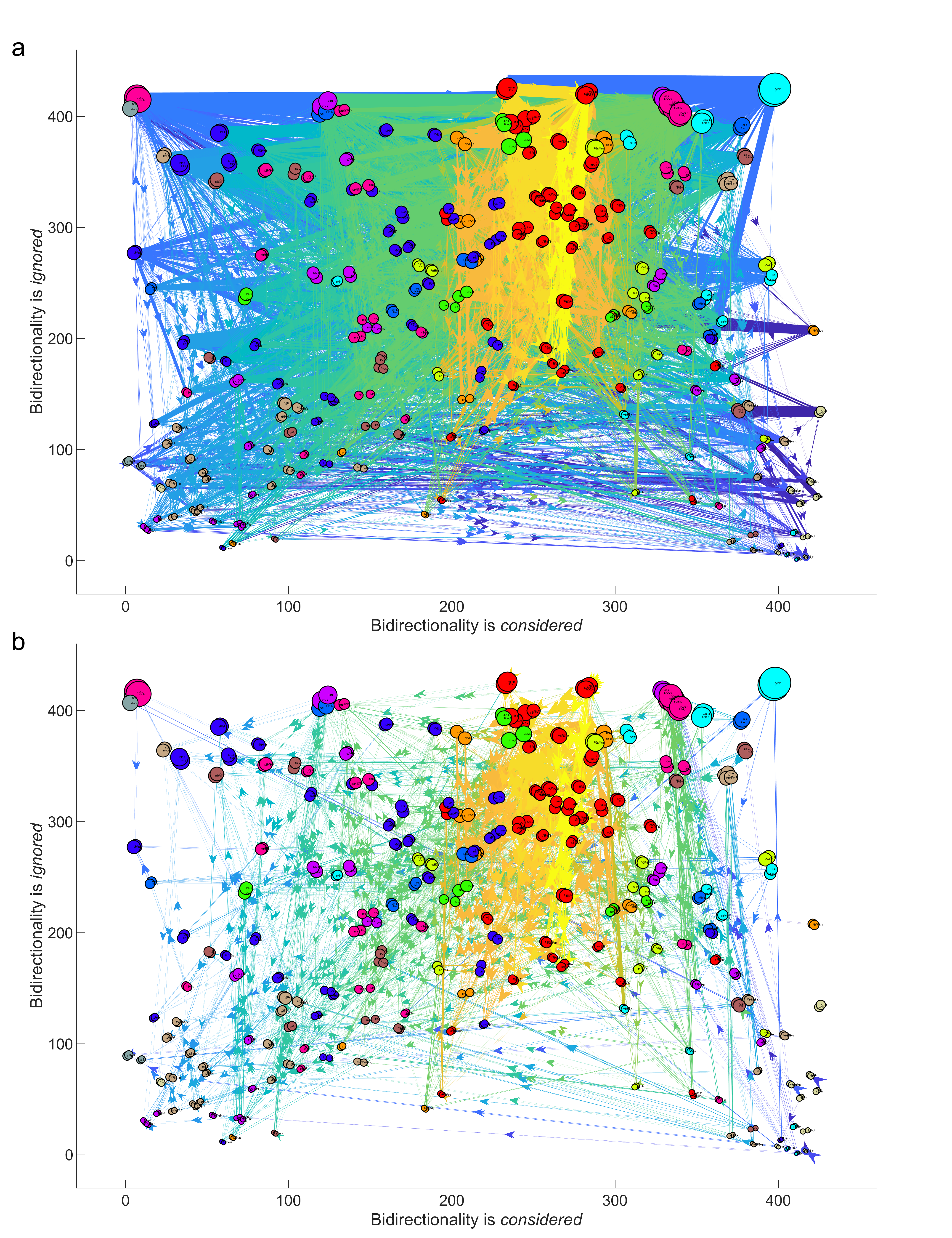

### Supplementary Figure 4

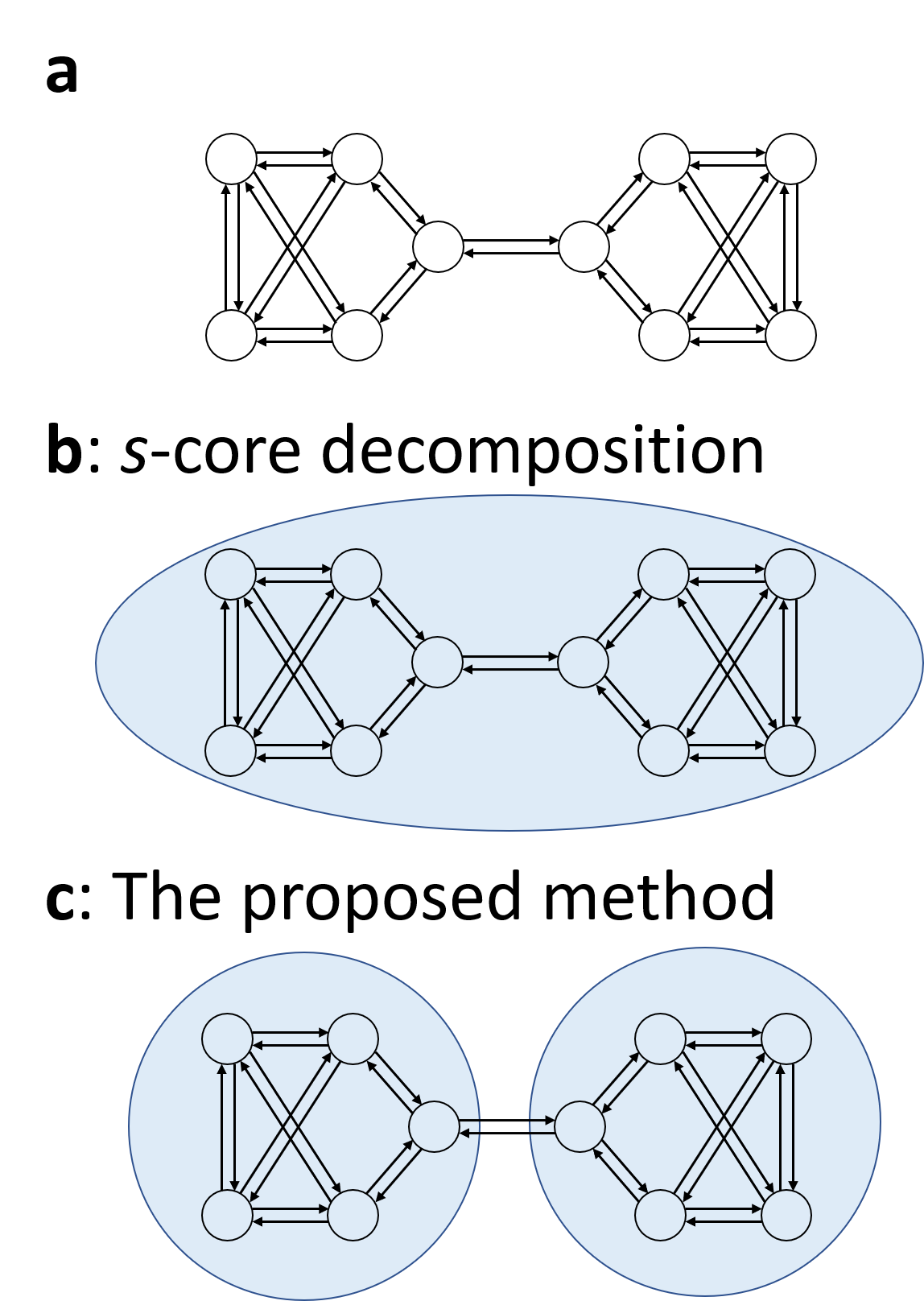
